# An applicable and efficient retrograde monosynaptic circuit mapping tool for larval zebrafish

**DOI:** 10.1101/2024.06.27.601104

**Authors:** Tian-Lun Chen, Qiu-Sui Deng, Kun-Zhang Lin, Xiu-Dan Zheng, Xin Wang, Yong-Wei Zhong, Xin-Yu Ning, Ying Li, Fu-Qiang Xu, Jiu-Lin Du, Xu-Fei Du

**Affiliations:** Institute of Neuroscience, Key Laboratory of Brain Cognition and Brain-Inspired Intelligence Technology, Center for Excellence in Brain Science and Intelligence Technology, Chinese Academy of Sciences, 320 Yue-Yang Road, Shanghai 200031, China; University of Chinese Academy of Sciences, 19A Yu-Quan Road, Beijing 100049, China; The Brain Cognition and Brain Disease Institute (BCBDI), Shenzhen Key Laboratory of Viral Vectors for Biomedicine, Shenzhen Institute of Advanced Technology, Chinese Academy of Sciences, Shenzhen, China; School of Life Science and Technology, ShanghaiTech University, 319 Yue-Yang Road, Shanghai 200031, China

## Abstract

The larval zebrafish is a vertebrate model for in vivo monitoring and manipulation of whole-brain neuronal activity. Tracing its neural circuits remains challenging. Here, we report an applicable methodology tailored for larval zebrafish to achieve efficient retrograde trans-monosynaptic tracing from genetically defined neurons via EnvA-pseudotyped glycoprotein-deleted rabies viruses. By combinatorially optimizing multiple factors involved, we identified the CVS strain trans-complemented with advanced expression of N2cG at 36 °C as the optimal combination. It yielded a tracing efficiency of up to 20 inputs per starter cell. Its low cytotoxicity enabled the viable labeling and calcium imaging of infected neurons 10 days post-infection, spanning larval ages commonly used for functional examination. Cre-dependent labeling was further developed to enable cell-type-specific input tracing and circuit reconstruction. We mapped cerebellar circuits and uncovered the ipsilateral preference and subtype specificity of granule cell-to-Purkinje cell connections. Our method offers an efficient way for tracing neural circuits in larval zebrafish.

## Introduction

Given the small transparent brain and accessible genetic manipulations, the larval zebrafish has been emerging as a promising animal model for systems neuroscience. Large-scale molecular, light- and electron-microscopical (LM and EM) morphological, and physiological datasets have been gathered over the past decade to decipher the identity and connectivity of its whole-brain cells (Kunst et al., 2019; Mu et al., 2012; Raj et al., 2020; Shainer et al., 2023; Svara et al., 2022; Yao et al., 2016; Zhang et al., 2021). Whole-brain neurons’ activity can be monitored and manipulated in awake as well as behaving zebrafish larvae, providing an unprecedented research paradigm for dissecting neural mechanisms of brain functions (Kappel et al., 2022; Loring et al., 2020; Mu et al., 2019; Naumann et al., 2016; Petrucco et al., 2023; Shang et al., 2024; Vanwalleghem et al., 2018). Further mechanistic insights into the roles of different cell types in brain-wide activity require effective tools for dissecting underlying neural circuits (Luo et al., 2018).

Virus-based tracing tools have opened new avenues for disentangling neural circuits (Nectow and Nestler, 2020; Xu et al., 2020). Among them, the EnvA-pseudotyped glycoprotein (G)-deleted rabies virus (RV) (RV*d*G[EnvA]) has achieved the most success (Miyamichi et al., 2011; Reardon et al., 2016; Wickersham et al., 2007). Its success derives from being a native retrograde transsynaptic tracer engineered for cell-type-specific targeting through the recognition between EnvA and its receptor TVA (Barnard et al., 2006) and crossing only one synaptic step via trans-complementation of rabies G (Miyamichi et al., 2011; Wickersham et al., 2007). This tool has revolutionized the structural analysis of neural circuits in rodents (Luo et al., 2018; Yao et al., 2023) and has continually evolved to adapt to functional studies (Chatterjee et al., 2018; Jin et al., 2024, 2023). However, its application in other species including zebrafish is retarded due to a lack of tailored methodologies.

The G-deleted RV (RV*d*G) infection of neurons in zebrafish was first demonstrated in 2009. Ten years later, RV*d*G[EnvA] was applied to trace input neurons in the adult zebrafish cerebellum, but the efficiency showed very low (Dohaku et al., 2019). Recently, the herpes simplex virus (HSV1) was used as a helper virus to deliver TVA and rabies G, leading to an increased efficiency of around one input per starter neuron in the brain of adult zebrafish (Satou et al., 2022). For larval zebrafish, efforts have been focused on the vesicular stomatitis virus (VSV), another rhabdovirus similar to RV but with anterograde spread (Kler et al., 2021; Ma et al., 2020; Mundell et al., 2015). However, these reported VSV tools for larval zebrafish appeared relatively high cytotoxicity and have not been iterated to utilize the EnvA-TVA system to achieve initial infection specificity (Kler et al., 2021; Ma et al., 2020; Mundell et al., 2015). Therefore, developing applicable efficient viral tracing methods is still an important mission in the field, and will boost the application of larval zebrafish in systems neuroscience research.

In the present study, we established applicable methodologies to implement RV*d*G[EnvA] for efficient retrograde trans-monosynaptic tracing from genetically defined neurons in larval zebrafish. We first demonstrated that the transient co-expression of the helper proteins TVA and rabies G in specific neurons through one-cell-stage microinjection of the GAL-UAS binary DNA plasmids could successfully help the RV*d*G[EnvA] microinjected later on to infect and spread from targeted starter cells. This simple two-round microinjection procedure ensures the high feasibility of our method. Then via practicing in the cerebellar circuit, we iteratively tested key factors that influence the tracing efficiency and cytotoxicity, which include the virus strain, G protein type, G expression level, and working temperature. We discovered an optimal combinatory condition: CVSdG trans-complemented with advanced expression of N2cG at 36 °C. The optimality lies in the high tracing efficiency and low cytotoxicity, embodied by a 20-fold increase in efficiency compared to previous studies in zebrafish (Dohaku et al., 2019; Satou et al., 2022) and a long enough time window (final fish age up to 2–3 weeks old) for conducting functional studies in larval zebrafish. Furthermore, by using Cre-expressing RV*d*G, we developed a transgenic reporter framework to enable input cell-type-specific tracing and circuit reconstruction based on single-neuron morphology. As an application demonstration, we applied the method to map the monosynaptic inputs to Purkinje cells (PCs) from granule cells (GCs) in the cerebellum and revealed the wiring relationship between morphological subtypes of GCs and PCs. These endeavors prove the applicability and efficiency of our method in dissecting interested circuits in larval zebrafish.

## Results

### Implementation of RV*d*G[EnvA]-based neural circuit tracing in larval zebrafish

To implement RV*d*G[EnvA]-based retrograde tracing in larval zebrafish, we first verified that larval zebrafish themselves do not express endogenous TVA. In wild type (WT) larvae without exogenous TVA expression, the injected SAD*d*G-mCherry[EnvA], a type of RV*d*G[EnvA], did not infect any cells (Figure 1—figure supplement 1A). The observed mCherry-positive puncta co-localized with microglia (Figure 1—figure supplement 1B), suggesting the clearance of uninfected viral particles by microglia. These data demonstrate that similar to mammals (Federspiel et al., 1994), zebrafish do not express endogenous TVA.

Then we specifically expressed TVA in neurons (Figures 1A and B) and found that injected SAD*d*G-mCherry[EnvA] infected TVA-expressing cells (Figure 1C and D, dashed yellow circles). The targeted expression of TVA and the fluorescent indicator enhanced green fluorescent protein (EGFP) in neurons was achieved through transient transgene of GAL4-UAS constructs in which the GAL4 activator is driven by *elavl3*, a pan-neuronal marker in zebrafish (Figure 1A, and “*UGT*” in Figure 1B). As the G protein is necessary for viral spread, SAD*d*G-mCherry[EnvA] did not spread beyond the initially infected neurons (i.e., starter cells) (Figure 1C and E–G).

**Figure 1.**
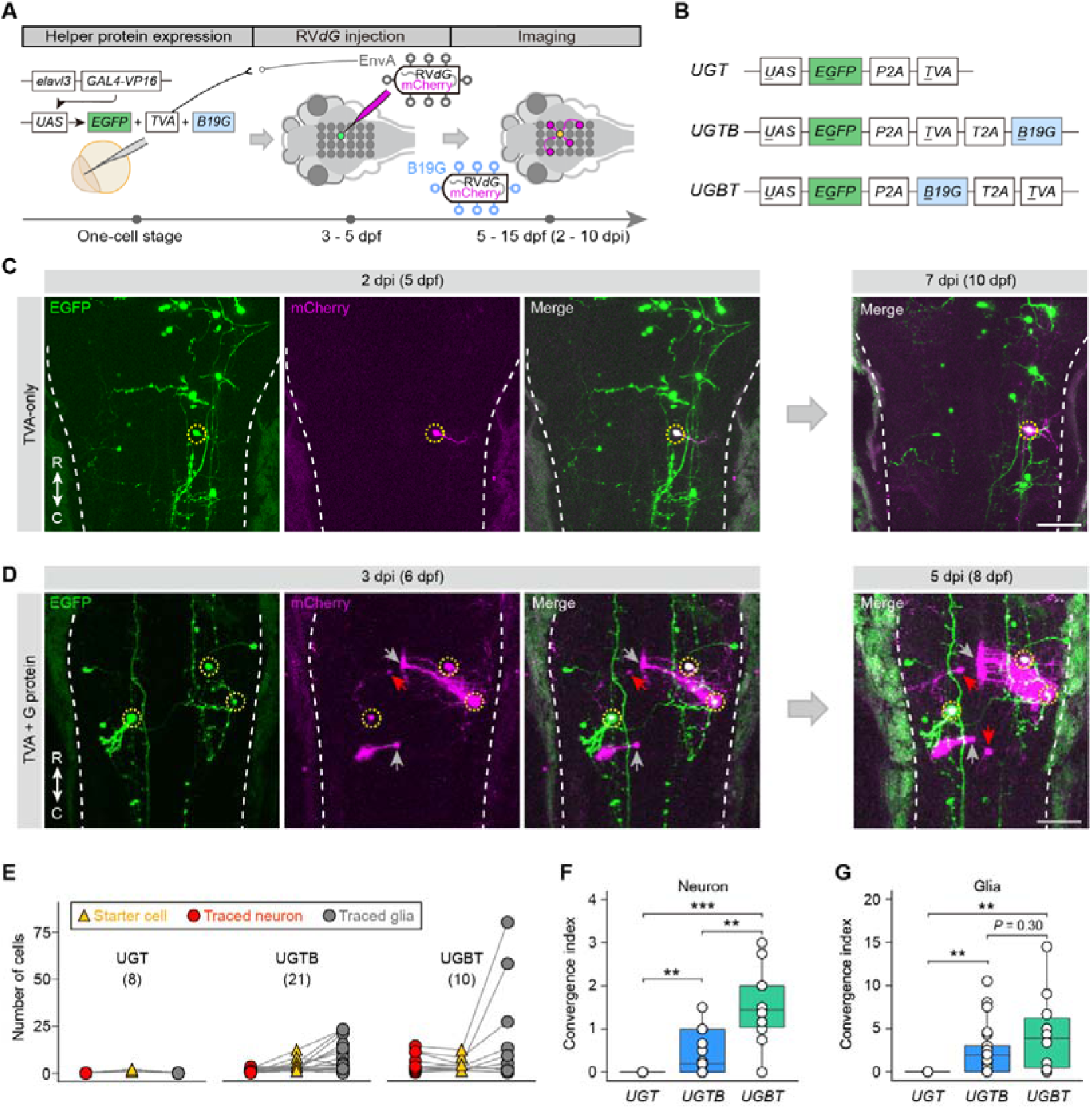
Implementation of RV*d*G[EnvA]-based neural circuit tracing in larval zebrafish. (A) Schematic of the method used to implement RV*d*G[EnvA]-based circuit tracing from genetically defined cell types in larval zebrafish. (B) Schematic of the three *UAS*-driven plasmids used to express the helper proteins TVA and G. (C and D) Time-lapse confocal images of the *nacre* larval hindbrain expressing EGFP and TVA using *UGT* (C), or EGFP, TVA, and B19G using *UGTB* (D) in randomly sparse neurons (with expression of *UAS* constructs driven by the *elavl3*:*GAL4-VP16* plasmid) and infected by SAD*d*G-mCherry[EnvA] (magenta) injected into the hindbrain. Dashed yellow circles, starter neurons (EGFP^+^/mCherry^+^); arrows, transneuronally labeled neurons (red) and radial glia (gray) (mCherry^+^/EGFP^−^); dashed white lines, hindbrain boundaries. C, caudal; R, rostral. Scale bars, 50 μm. (E) Plots of the numbers of starter neurons, transneuronally traced neurons and glia in each larva expressing one of the three helper plasmids shown in (B) and injected with SAD*d*G-mCherry[EnvA]. Virus injections were carried out at 3–5 dpf; exact ages for individual animals are listed in Figure 1—table supplement 2. Data were collected between 6 and 10 dpi. Numbers in brackets indicate the number of larvae examined. (F and G) Boxplots of the convergence index for the connection of transneuronally traced neurons (F) and glia (G) to starter neurons shown in (E). Center, median; bounds of box, first and third quartiles; whiskers, minimum and maximum values. \*\**P* < 0.01, \*\*\**P* < 0.001 (nonparametric two-tailed Mann-Whitney test).

We next examined whether viral particles could spread to other cells by complementing the G protein in starter neurons. As the efficiency of virus spread is reported to be positively correlated with the level of G expression in starter neurons (Callaway and Luo, 2015; Miyamichi et al., 2013), we generated two helper plasmids, *<u>U</u>AS:E<u>G</u>FP-P2A-<u>T</u>VA-T2A-<u>B</u>19G* (*UGTB*) and *<u>U</u>AS:E<u>G</u>FP-P2A-<u>B</u>19G-T2A-<u>T</u>VA* (*UGBT*), to search for a plasmid design with higher G expression while ensuring the co-expression of TVA and G (Figure 1B, see Materials and methods). The position of the RV G-protein-coding gene SAD B19G was varied in the tri-cistronic 2A constructs for adjusting its expression level (Liu et al., 2017). As shown in Figure 1D, besides starter cells (EGFP^+^/mCherry^+^, dashed yellow circles) and uninfected TVA-expressing cells (EGFP^+^ only), we observed mCherry^+^ only cells (arrows) of which the number increased over time, indicating the virus spread beyond the starter cells by trans-complementation with G.

Notably, the traced cells included both neurons and radial astroglia (hereafter referred to as glia) identified by their distinct morphology (Figure 1D, red and gray arrows, respectively). Trans-infection of glia has also been observed in mice (Beier, 2021; Marshel et al., 2010), and its mechanism remains unknown. Glial labeling was frequently observed in close proximity to the soma and dendrites of starter neurons (18 out of 21 cases, Figure 1D), a pattern that was further confirmed in an independent set of experiments using *vglut2a* to define starter neurons (14 out of 17 cases; Figure 1—figure supplement 2). These observations suggest that transneuronally labeled glial cells are closely associated with starter neurons, potentially through synaptic or perisynaptic interactions.

We also expressed TVA and G in glial cells using the *gfap* promoter and performed rabies tracing with SAD*d*G-mCherry[EnvA]. Under this condition, we observed robust initial infection of TVA-positive glia, confirming TVA-dependent, cell-type-specific viral entry. No labeling of nearby TVA-negative neurons or glial cells was detected (0 out of 3 starters from 3 larvae; Figure 1,2—figure supplement 1A), indicating an absence of detectable trans-cellular spread from glial starter cells. By contrast, different outcomes were observed when alternative rabies strain and glycoprotein were used (see below).

Convergence index (CI), defined as the number of trans-infected cells divided by that of starter neurons, was then used to estimate the tracing efficiency (Callaway and Luo, 2015; Miyamichi et al., 2011). We calculated the CI of traced neurons and glia separately, and found a higher tracing efficiency in larvae expressing *UGBT* than those expressing *UGTB* (Figure 1E–G and Figure 1,2—table supplement 1 and 2; CI_neuron_: 1.55 ± 0.29 *vs*. 0.44 ± 0.11, *P* < 0.01; CI_glia_: 4.41 ± 1.48 *vs*. 2.50 ± 0.65, *P* = 0.30; mean ± SEM), suggesting that *UGBT* is better for G expression and tracing efficiency. Taken together, these results demonstrate that in larval zebrafish, transneuronal tracing from genetically defined cell types can be implemented by combining the injection of RV*d*G[EnvA] with the co-expression of helper proteins mediated by the GAL4-UAS system.

## Optimization and verification of the retrograde monosynaptic tracing

To optimize this system for high tracing efficiency and application feasibility in circuit research, we iterated it step-by-step in the cerebellum via testing different combinations of RV strains (SAD-B19 *vs*. CVS-N2c), G proteins (SAD B19G *vs*. CVS N2cG), and rearing temperatures (28 °C *vs*. 36 °C) (Figure 2A). The cerebellar structure is evolutionarily conserved in zebrafish compared with mammals (Bae et al., 2009; Hibi and Shimizu, 2012). In zebrafish, Purkinje cells (PCs), the central cell type in the cerebellar circuit, receive excitatory inputs from parallel fibers and climbing fibers originated respectively from granule cells (GCs) and inferior olivary cells (IOCs), as well as local inhibitory inputs from stellate cells (SCs) (Bae et al., 2009; Hibi and Shimizu, 2012) (Figure 2B). The helper proteins G and TVA were co-expressed in PCs by using the plasmid in *UGBT* mode, driven by *cpce*, a *ca8* promoter-derived PC-specific enhancer element (Bae et al., 2009; Namikawa et al., 2019) (Figure 2A).

**Figure 2.**
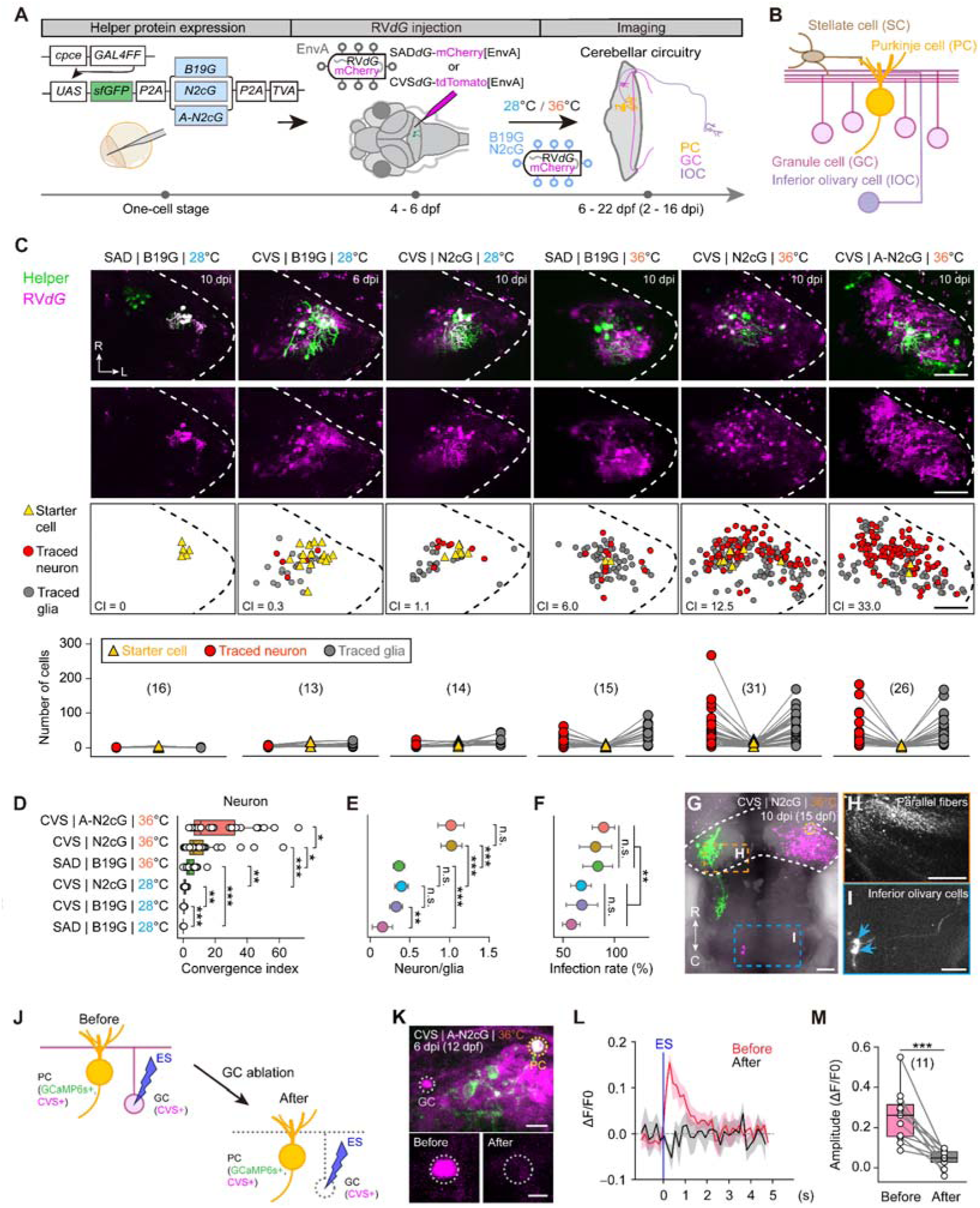
Optimization and verification of RV*d*G[EnvA]-based retrograde monosynaptic tracing. (A) Schematic of the method used to trace inputs to PCs with RV*d*G[EnvA] in larval zebrafish. The *cpce:GAL4FF* plasmid and one of the three *UAS* helper plasmids were injected into *nacre* fish and combined with subsequent injection of SAD*d*G-mCherry[EnvA] or CVS*d*G-tdTomato[EnvA]. The virus-injected larvae were raised at either 28 or 36 °C, and the resulting labeling was monitored from 2 to 16 dpi. (B) Schematic of the zebrafish cerebellar circuit related to inputs to PCs. (C) Top, examples of in situ complementation in PCs under six tracing conditions. Confocal images and corresponding schematic illustrations of one cerebellar hemisphere are shown. Bottom,plots of the numbers of starter neurons, transneuronally traced neurons and glia in each larva under one of the six tracing conditions at 6–10 dpi. Virus injections were carried out at 4–6 dpf; exact ages for individual animals are listed in Figure 1,2—table supplement 2. Dashed lines, the cerebellar boundaries. CI, convergence index of traced neurons. Numbers in brackets indicate the number of larvae examined. (D–F) CI for the connection of transneuronally traced neurons to starter PCs (D), the ratio of the number of traced neurons to that of glia (E), and the proportion of infected fish (F, 3 to 5 independent experiments) under the six tracing conditions shown in (C, bottom). (G) Confocal image of a larva infected with CVS*d*G-tdTomato[EnvA] that was trans-complemented with CVS N2cG at 36 °C. Dashed yellow circle, starter PC; dashed white line, the cerebellar boundary. (H and I) Enlarged view of the boxed regions in (G), showing transneuronally traced parallel fibers of GCs (H) and two IOCs (I). Blue arrows, somata of IOCs. (J) Schematic of the experiment for verifying the functional connection between traced GCs and the starter PC in *nacre* larvae injected with *cpce:GAL4FF* and advanced *UG6sNT-A* plasmids and infected with CVS*d*G-tdTomato[EnvA] at 36 °C. Ca^2+^ imaging was conducted on the starter PC in response to electrical stimulation (ES) at the GC position before and after two-photon laser ablation of the GC .(K) Representative image (top) showing both the stimulated GC and the simultaneously recorded PC. Bottom panels display the stimulated GC before (left) and after (right) two-photon laser ablation. Dashed yellow circle, recorded PC (GCaMP6s^+^/tdTomato^+^); dashed white circles, stimulated GC (GCaMP6s^−^/tdTomato^+^). (L) Average Ca^2+^ response (5 trials) of the same starter PC before (red) and after (black) GC ablation. The shadowed area represents SEM. (M) Quantification of the amplitude of PC Ca^2+^ responses before and after GC ablation. The number in bracket indicates the number of GC-PC pairs examined. Scale bars, 50 μm (C, G–I), 10 μm (K, top), 5 μm (K, bottom). SAD/CVS|B19G/N2cG/A-N2cG|28/36 °C, the tracing conditions written as the G-deleted RV strain that was injected | the helper glycoprotein used for trans-complementation | fish rearing temperature after virus injection. C, caudal; L, lateral; R, rostral. Boxplots in (D and M): center, median; bounds of box, first and third quartiles; whiskers, minimum and maximum values. n.s., not-significant; \**P* < 0.05, \*\**P* < 0.01, \*\*\**P* < 0.001 (nonparametric two-tailed Mann-Whitney test in (D–F); two-tailed unpaired Student’s *t* test in (L)). Error bars denote SEM.

With the same G protein SAD B19G and rearing temperature at 28 °C, CVS*d*G[EnvA] exhibited around a 6-fold increase in neuronal tracing efficiency compared to SAD*d*G[EnvA] (Figure 2C and D, Figure 2—figure supplement 1A–C, and Figure 1,2—table supplement 1 and 2; CI_neuron_: 0.41 ± 0.07 *vs*. 0.06 ± 0.03, *P* < 0.001). After replacing SAD B19G with CVS N2cG (*UGNT*, see Materials and methods) to complement CVS*d*G[EnvA], the tracing efficiency was further increased by ∼2 times (Figure 2C and D, and Figure 1,2—table supplement 1 and 2; CI_neuron_: 1.02 ± 0.18 *vs*. 0.41 ± 0.07, *P* < 0.01). In addition, CVS*d*G[EnvA] showed a greater tendency to spread to neurons rather than glia in comparison with SAD*d*G[EnvA] (Figure 2C and E). In all experiments, virus injection was performed at 4–6 days post-fertilization (dpf). The labeling of input neurons usually emerged around 2 days post injection (dpi), and all CIs were calculated at 6–10 dpi, during which the viral transfer was relatively stable (Figure 2—figure supplement 1C).

To further assess the performance of CVS rabies strains complemented with the N2cG glycoprotein, we applied this tracing strategy to glial starter cells. Consistent with the SAD condition, no neuronal labeling was observed. However, labeling of nearby TVA-negative glial cells was detected, indicating glia-to-glia transmission (3 out of 6 starters from 6 larvae; Figure 1,2—figure supplement 1B). This is consistent with previous rabies tracing studies reporting astrocyte-associated viral spread, including astrocyte–astrocyte interactions (Clark et al., 2021).

After elevating the temperature from 28 °C to 36 °C for rearing virus-injected larvae, the neuron tracing efficiency of SAD*d*G[EnvA] became obvious (Figure 2C and D, Figure 2—figure supplement 1D, and Figure 1,2—table supplement 1 and 2; CI_neuron_: 4.96 ± 1.01 *vs*. 0.06 ± 0.03, *P* < 0.001); for CVS*d*G[EnvA] trans-complemented with N2cG, the efficiency further increased by one order of magnitude (Figure 2C and D, Figure 2—figure supplement 1E, and Figure 1,2—table supplement 1 and 2; CI_neuron_: 11.61 ± 2.43 *vs*. 1.02 ± 0.18, *P* < 0.001). In addition, the elevated temperature also significantly increased the proportion of initially infected larvae (Figure 2F) and the tendency of viral spread to neurons (Figure 2E). This efficient trans-synaptic viral transfer from PCs allowed clear labeling of GC parallel fibers and long-range contralateral inputs from IOCs (Figure 2G–I). We then generated a *Tetoff* element-based advanced helper plasmid (*UGNT-A*) to enhance the expression level of CVS N2cG (A-N2cG) (Figure 2—figure supplement 1F, see Materials and methods), and this optimization resulted in a nearly two-fold increase in the neuron tracing efficiency (Figure 2C and D, Figures 2—figure supplement 1F and G, and Figure 1,2—table supplement 1 and 2; CI_neuron_: 20.35 ± 3.45 *vs*. 11.61 ± 2.43, *P* < 0.05). The improvement was also observed for tracing glia (Figure 2C, Figure 2—figure supplement 1C, G, H and I, and Figure 1,2—table supplement 1 and 2).

Time-lapse analysis of neuronal and glial labeling showed that both cell types accumulated over time, but glial labeling markedly slowed between 6 and 10 dpi (Figure 2—figure supplement 1C and G). We therefore divided the timeline into early (2–6 dpi) and late (6–10 dpi) phases and quantified the daily change in mean CI as the labeling rate (R). A labeling rate index (R_glia_ − R_neuron_) was used to compare glial and neuronal labeling dynamics. The index was positive in the early phase (0.87 ± 0.43) but became negative in the late phase (−0.80 ± 0.39), indicating that neuronal labeling outpaced glial labeling over time (Figure 2—figure supplement 2).

To investigate the basis of this slowdown, we used the time-lapse data to examine the persistence of labeled glia and found that glial fluorescence progressively diminished after transneuronal labeling without apoptotic debris, consistent with abortive infection (Tian et al., 2018). Supporting this interpretation, we quantified the proportion of glial cells labeled at 2 dpi and 4dpi that retained fluorescence over time and found that labeling largely diminished by 6 dpi (∼11 dpf) in both groups (Figure 2—figure supplement 3), suggesting that the decline is more closely associated with larval age than with infection duration. Despite this loss, the convergence index continued to increase, implying that ongoing de novo glial infection outpaces the loss of labeled glia.

To further functionally test the synaptic specificity of RV retrograde spread in the cerebellar circuit, we conducted in vivo Ca^2+^ imaging of starter PCs while simultaneously activating single traced GCs via electrical stimulation (Figure 2J, see Materials and methods). To monitor the activity of starter PCs, the fluorescent protein in the helper plasmid was replaced with the Ca^2+^ indicator GCaMP6s (*UG6sNT-A*, see Materials and methods). In total, we stimulated 33 individual GCs across 32 animals, among which 15 elicited detectable calcium responses in putative postsynaptic PCs. In 11 of these 15 cases where the stimulated GC was successfully ablated, the GC-evoked Ca^2+^ activity in starter PCs was abolished after two-photon laser-based ablation of the GC (Figure 2K–M and Figure 1,2—table supplement 2). These results indicate that RV can retrogradely spread to GCs which are functionally connected with starter PCs, confirming the capability of RV to trace monosynaptic inputs in larval zebrafish.

### Time window for normal neuronal health after RV infection

In sum, we have identified the optimal conditions for efficient retrograde monosynaptic tracing in zebrafish larvae: CVS*d*G[EnvA] trans-complemented with A-N2cG via the GAL4-UAS system at 36 °C. The high efficiency of these optimal tracing conditions prompted concerns regarding the potential cytotoxicity of CVS*d*G and CVS N2cG. In line with previous reports in mice (Reardon et al., 2016), we found lower toxicity of CVS*d*G compared with SAD*d*G. At 36 °C, the survival rate of larvae infected with CVS*d*G was significantly higher than those infected with SAD*d*G (Figure 3A; 75% ± 3% *vs*. 52% ± 7%, *P* < 0.05). Furthermore, by using the time-lapse imaging as shown in Figure S2, we often observed that early emerged starter cells disappeared in larvae infected with SAD*d*G (Figure 3B). As new starter cells continued to appear from 2 to 6 dpi (Figure 3C), we utilized starter cells that emerged at 2 dpi to quantify their lifetime. Consistently, starter cells infected by CVS*d*G survived longer than those infected by SAD*d*G (Figure 3D; 10.0 ± 0 days *vs*. 7.5 ± 0.48 days, *P* < 0.001). Notably, the viability of starter cells decreased when CVS*d*G was trans-complemented with A-N2cG (Figure 3D; 10.0 ± 0 days *vs*. 8.7 ± 0.48 days, *P* < 0.001).

**Figure 3.**
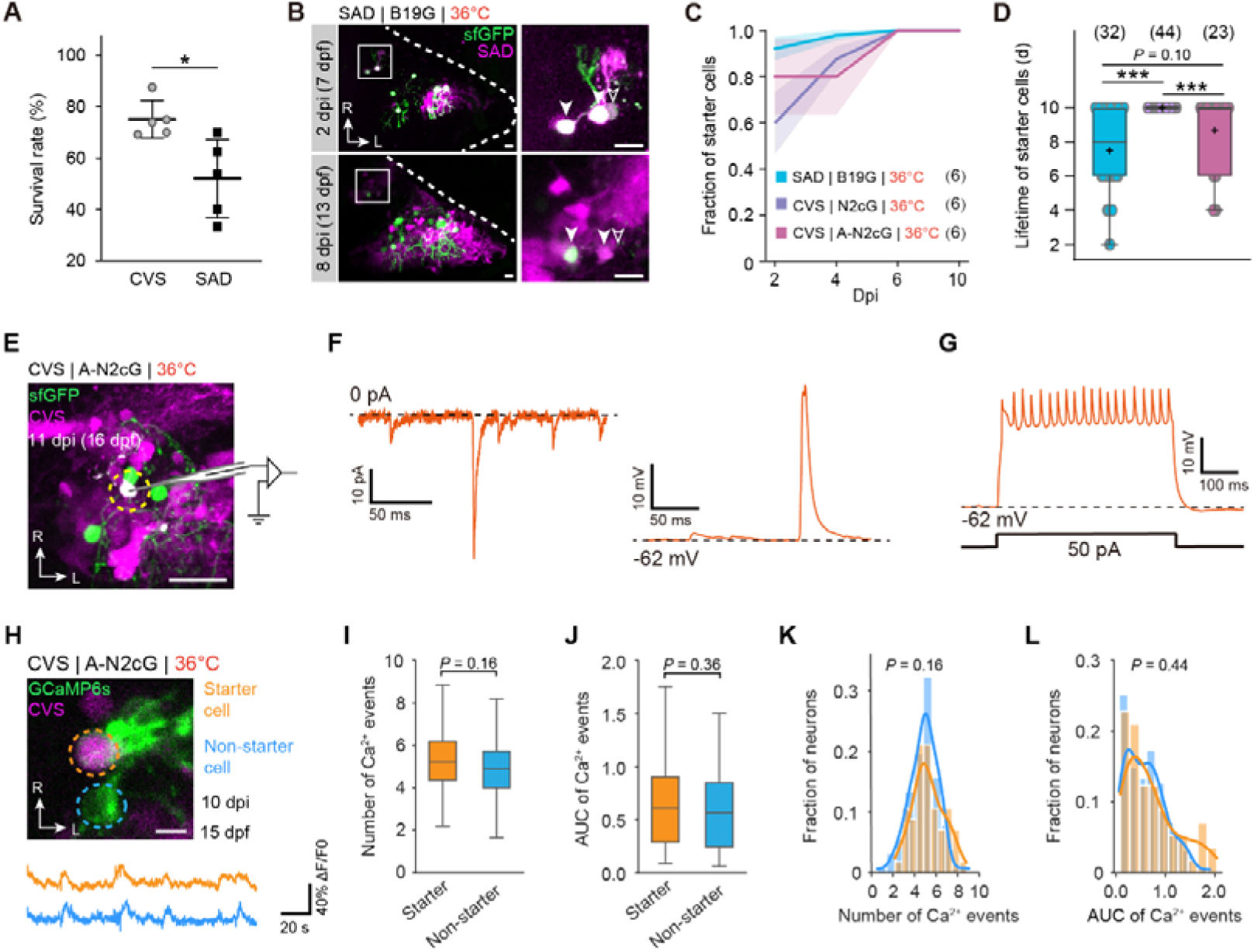
Neuronal health and physiology appeared normal for at least 10 days after RV infection. (A) Survival rate of larvae injected with SAD*d*G-mCherry[EnvA] or CVS*d*G-tdTomato[EnvA]. Five independent experiments were repeated. (B) Time-lapse confocal images showing that a starter cell (unfilled white arrowhead) present at 2 dpi disappears by 8 dpi, and two starter cells (white arrowheads) remain present from 2 to 8 dpi. Boxed regions are enlarged on the right. Dashed lines indicate the cerebellar boundaries. (C) Cumulative proportion of starter cells that have appeared at 2, 4, 6, and 10 dpi to the total number of observed starter cells under the three tracing conditions, showing the continuous appearing of starter cells at 2–6 dpi. Numbers in brackets indicate the number of larvae examined. The shadowed area represents SEM. (D) Summary of the lifetime of starter cells appeared at 2 dpi under the three tracing conditions. The color scheme follows the same conventions shown in (C). Numbers in brackets indicate the number of starter cells examined. (E) Schematic illustration of in vivo whole-cell recording on a starter PC at 11 dpi. (F) Example traces of spontaneous EPSCs (left) and EPSPs (right) from starter PCs. (G) Example trace of spikes in response to current-step injection into a starter PC. (H) Confocal image (top) and Ca^2+^ responses (bottom) of a starter PC (dashed orange circle) and non-starter PC (dashed blue circle) to visual stimuli at 10 dpi. (I–L) Boxplots (I and J) and distributions (K and L) of Ca^2+^ activity event number (I and K) and area under curve (AUC) of Ca^2+^ activity events (J and L) of starter PCs (n = 88) and non-starter PCs (n = 127) in larvae (N = 8) at 6–10 dpi. All tracing experimental procedures were identical to those described in Figure 2. In panels H–L, the helper plasmid *UG6sNT-A* was used to express GCaMP6s for calcium imaging of PCs. Scale bars, 5 μm (H), 10 μm (B), 20 μm (E). SAD/CVS|B19G/N2cG/A-N2cG|28/36 °C, the tracing conditions written as the G-deleted RV strain that was injected | the helper glycoprotein used for trans-complementation | fish rearing temperature after virus injection. L, lateral; R, rostral. Boxplots in (D, I, and J): center, median; cross symbol, mean; bounds of box, first and third quartiles; whiskers, minimum and maximum values. n.s., not-significant; \**P* < 0.05; \*\*\**P* < 0.001 (two-tailed unpaired Student’s *t* test). Error bars denote SEM.

To further test whether the physiological functions of starter cells were impaired by CVS*d*G trans-complemented with A-N2cG, we conducted in vivo whole-cell recordings on starter PCs at around 10 dpi (Figure 3E, see Materials and methods). Infected PCs exhibited normal spontaneous excitatory postsynaptic currents and potentials (sEPSCs and sEPSPs), and current injection-evoked firing patterns (Figures 3F and G) (Sengupta and Thirumalai, 2015). Then we examined Ca^2+^ activity of infected (i.e., starter cells) and uninfected PCs (i.e., non-starter cells) in response to complex visual stimulation (Knogler et al., 2019) (see Materials and methods). No significant differences were detected between starter and non-starter PCs (Figure 3H–L). Taken together, these results indicate that under the optimal tracing conditions, infected neurons can remain relatively healthy up to 10 days after infection in larval zebrafish, allowing for functional probing of traced neural circuits.

### Cell-type-specific input tracing and circuit reconstruction in the cerebellum

One important application of neural circuit tracing is to map the structural connectivity of specific neuron types to build the brain connectome atlas. As shown above, even under the optimal tracing conditions, glia still constituted around 47% on average of the traced cells (see Figure 2C and E). This will certainly affect the discrimination of traced neurons and their neural fibers. To confine retrograde tracing to neurons instead of glia, we generated a Cre-Switch (Saunders et al., 2012) transgenic reporter fish *Tg(elavl3:Tetoff-DO_DIO-Hsa.H2B-mTagBFP2_tdTomato-CAAX)*. This reporter expresses pan-neuronally nuclear-localized blue fluorescent protein (mTagBFP2) by default, which serves as a reference for image registration and data integration. It also achieves Cre-dependent neuron-specific labeling with tdTomato-CAAX via the *elavl3* promoter (Figure 4A, see Materials and methods). We employed this reporter together with Cre-expressing CVS*d*G (CVS*d*G-Cre[EnvA]) to map neuronal inputs to PCs in the cerebellum (Figure 4B). The transgenic labeling of traced neurons showed much brighter and sharper fluorescence compared with virus-assisted labeling (Figure 4C). Among all 13 fish examined, we detected tdTomato-CAAX signals exclusively in neurons without any labeling of glia (Figure 4C), facilitating the reconstruction of traced neural circuits (Figure 4D). We noticed a relatively lower tracing efficiency of this Cre-dependent tracing system (Figures 4C and 2C; CI_neuron_: 5.20 ± 1.09 *vs*. 11.61 ± 2.43). Efforts are underway to enhance the completeness of transgene expression and improve the efficiency of Cre-mediated recombination.

**Figure 4.**
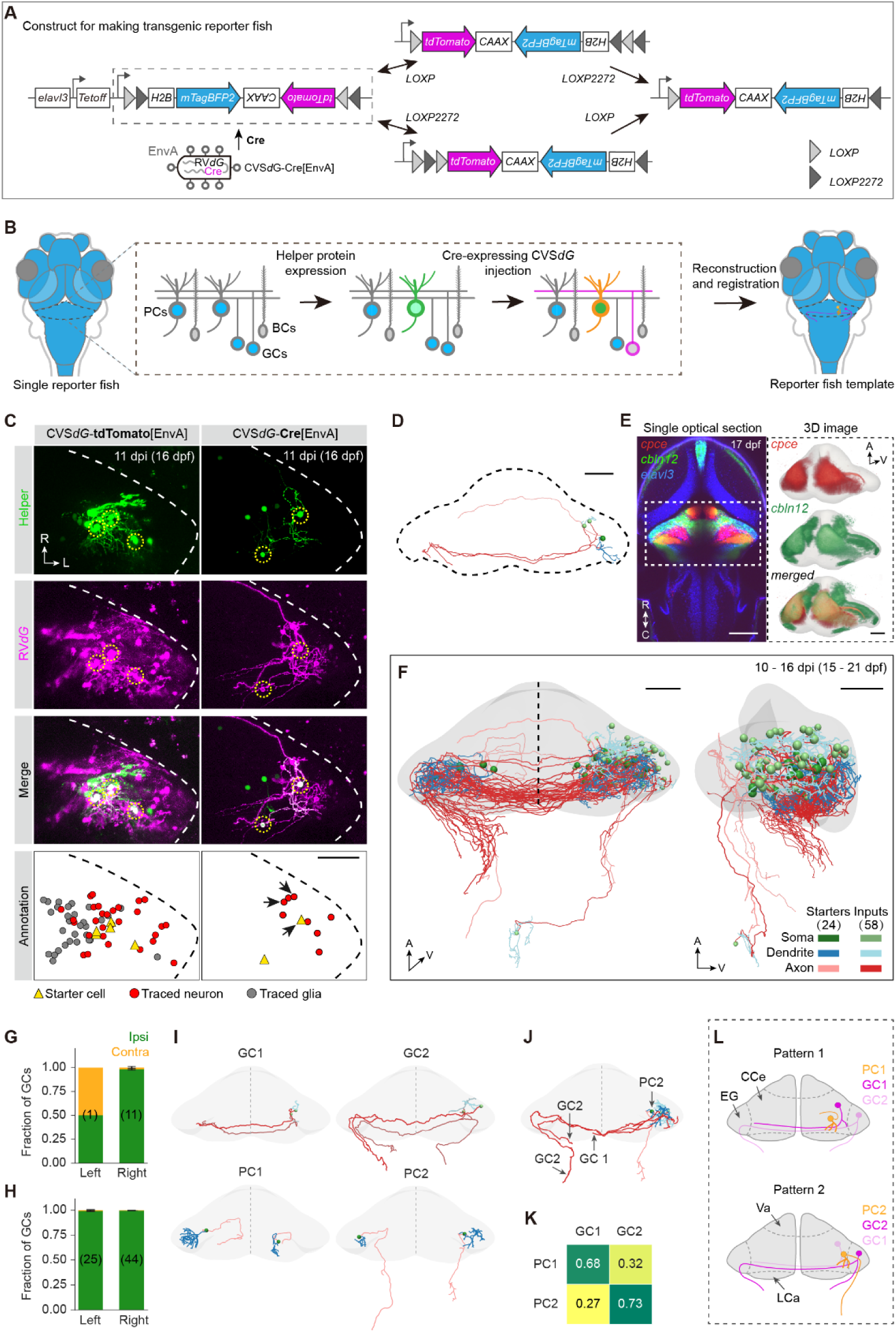
Cell-type-specific input tracing and circuit reconstruction in the cerebellum. (A) A single Cre-Switch (Saunders et al., 2012) transgene (*elavl3:Tetoff-DO_DIO-Hsa.H2B-mTagBFP2_tdTomato-CAAX*; abbr., *elavl3DoDioBR*) was generated to achieve a reference transgene (*elavl3:H2B-mTagBFP2*) for image registration and the Cre-dependent reporter expression (*tdTomato-CAAX*) specifically in neurons (*elavl3*^+^). Two ORFs are flanked by two pairs of LOX sites in opposite orientations, with activation of the forward-oriented *H2B-mTagBFP2* ORF under the *elavl3* promoter in global neurons. After Cre-expressing virus infection, Cre recombines either pair of LOX sites (LOXP (gray triangles) or LOXP2272 (black triangles)), resulting in stable inversion of the two ORFs and activation of the *tdTomato-CAAX* ORF under the *elavl3* promoter in initially infected starter neurons and transneuronally infected input neurons. (B) Schematic of building PC input atlas in larval zebrafish. The *elavl3DoDioBR* reporter fish (pan-neuronal H2B-mTagBFP2^+^) expressing helper proteins in PCs (sfGFP^+^/mTagBFP2^+^) were injected with Cre-expressing CVS*d*G[EnvA], which were trans-complemented in starter PCs and spread to direct inputs, resulting in Cre-dependent expression of the reporter protein (tdTomato-CAAX) in *elavl3*^+^ starter PCs (sfGFP^+^/tdTomato^+^/mTagBFP2^−^) and input GCs (sfGFP^−^/tdTomato^+^/mTagBFP2^−^). The pan-neuronally-expressing *H2B-mTagBFP2* in *elavl3DoDioBR* serves as the reference and bridge for the registration of reconstructed circuits. BCs, Bergmann glia; GCs, granule cells; PCs, Purkinje cells. (C) Example of in situ complementation (11 dpi) in *nacre* larvae (left) and *elavl3DoDioBR* reporter larvae (right). Both were co-injected with *cpce-GAL4FF* and *UGNT* plasmids, and infected with CVS*d*G-tdTomato[EnvA] and CVS*d*G-Cre[EnvA], respectively. All larvae were maintained at 36 °C. In the reporter larvae, Cre-dependent tdTomato expression was confined to neurons, driven by neuron-specific *elavl3* promoter. Confocal images (top) and corresponding schematic illustrations (bottom) are shown. Dashed yellow circles, starter PCs; dashed lines, the cerebellar boundaries. Black arrows indicate the reconstructed neurons in (D). (D) Dorsal 3D view of the reconstructed starter PC and its two input GCs. Dashed line indicates the cerebellar boundary. The color scheme follows the same conventions as in (F). (E) Left, single horizontal section view of *Tg(2×en.cpce-E1B:tdTomato-CAAX)* (red), *Tg(cbln12:GAL4FF);Tg(5×UAS:EGFP)* (green), and *elavl3DoDioBR* (blue) templates aligned in a common coordinate space. Right, volume rending of the cerebellar area (dashed white rectangle on the left) of the templates labeling PCs (red) and GCs (green) in cerebellum region outline (gray). (F) Dorsal (left) and lateral (right) 3D view of reconstructed starter PCs (n = 24) and input GCs (n = 58) in the cerebellum (gray). Different subcellular parts of PCs and GCs are color-coded. Data were collected from 13 larvae at 10–16 dpi (15–21 dpf). (G and H) Ipsilateral *vs*. contralateral proportions of GC inputs to PCs in the left and right cerebellum. Analysis was performed using the reconstructed data in (E) and the viral tracing data shown in Figure 2C,D for (G) and (H), respectively. Numbers in brackets indicate the number of larvae examined which only had starter PCs on one cerebellar hemisphere. (I) Subtypes of reconstructed PCs and GCs. Two typical cells are shown for each subtype. (J) Example showing three GCs traced from a single PC. (K) Summed input proportion of two subtypes of GCs to two subtypes of PCs. Cases with only one subtype of PC and clearly identifiable GCs were included. The cell numbers are as follows: PC1, 12; PC2, 6; GC1, 19; GC2, 18. (L) Schematic representation of subtype-specific connectivity patterns from GCs to PCs. High saturation of color indicates a higher input strength. CCe, corpus cerebelli; EG, eminentia granularis; LCa, lobus caudalis cerebelli; Va, valvula cerebelli. Scale bars, 50 μm. A, anterior; C, caudal; L, lateral; R, rostral; V, ventral. Dashed vertical lines indicate the midline of the cerebellum.

To demonstrate the feasibility of our tracing system in building and analyzing brain input maps, we then generated a three-dimensional (3D) reference template brain from the transgenic reporter fish at 17 dpf (see Materials and methods). To incorporate cerebellar expression patterns, we generated the transgenic lines *Tg(2×en.cpce:tdTomato-CAAX)* and *Tg(cbln12:GAL4FF)* to label PCs and GCs, respectively. By utilizing H2B-mTagBFP2 expression in the reporter fish as a bridge, we mapped the two cerebellar expression patterns and circuit tracing results of 13 fish mentioned above onto the common coordinate space of the reference template (Figure 4B and E, see Materials and methods). We delineated the cerebellar brain area and reconstructed each starter PC (n = 24) and its input GCs (n = 58) with clearly visible morphology (Figure 4E and F), resulting in a digital cellular-resolution input atlas of cerebellar PCs (Figure 4F).

This input atlas clearly shows an ipsilateral preference of afferents from GCs to PCs (Figure 4G), which was in accordance with the summarized results of non-Cre-dependent viral tracing data (Figures 4H, and 2C and D). Interestingly, all reconstructed GCs also innervated contralaterally by crossing the midline. We found two morphological subtypes for both GCs and PCs in the reconstruction, and each included one local subtype with axonal projections within the cerebellar area and one long projection subtype with axons extending caudally to the dorsal hindbrain (Figure 4I). This classification of GC subtypes (GC1 and GC2) is consistent with previous studies (Bae et al., 2009; Matsui et al., 2014; Takeuchi et al., 2015): GC1 corresponds to rostromedial GCs with T-shaped axons and somata located in the valvula cerebelli (Va) and corpus cerebelli (CCe), whereas GC2 corresponds to caudolateral GCs that project to the dorsal hindbrain and innervate crest cells, with somata in the eminentia granularis (EG) and lobus caudalis cerebelli (LCa). Interestingly, although a single PC can receive inputs from the two subtypes of GCs (Figure 4J), the two subtypes of PCs prefer inputs from different subtypes of GCs (Figure 4K and L). These results demonstrate how cellular-level connectivity can advance our understanding of circuit wiring patterns.

Together, the development of the Cre-dependent tracing tool empowers the RV*d*G-based circuit tracing method, enabling input-specific tracing, cellular-level circuit reconstruction, and integrated mapping and analysis of afferent connections in larval zebrafish.

## Discussion

Understanding brain functions requires systematic dissection of the involved neural circuits. Here, we developed a highly feasible and efficient implementation of the viral tracing tool RV*d*G[EnvA] for circuit studies in larval zebrafish (Figure 1–4—table supplement 1). Through step-by-step improvement, we identified an optimal tracing condition for larval zebrafish, which includes the CVS*d*G strain, the native G protein of the CVS (i.e., N2cG), an enhanced expression system for the G protein, and elevated rearing temperature. Our method exhibited a 20-fold increase in tracing efficiency compared to previous studies in adult zebrafish (Dohaku et al., 2019; Satou et al., 2022), and demonstrated low cytotoxicity. Moreover, we established a custom-designed transgenic reporter framework, which enables genetic access to specific input cell types and allows for cellular-resolution reconstruction and alignment of traced circuits. We demonstrated the application of this framework to decipher cell-type specific connections from GCs to PCs in the cerebellum.

### Helper protein expression design that guarantees high feasibility of the tracing method

Precise spatial control of helper protein expression (i.e., TVA and G) in starter cells is critical for the application of RV*d*G[EnvA]-based circuit tracing. In zebrafish, previous studies expressed these proteins through generating transgenic fish lines (Dohaku et al., 2019; Satou et al., 2022) or injecting helper viral vectors (Kler et al., 2021; Ma et al., 2020; Satou et al., 2022). Both strategies allow genetic targeting of specific neuronal types or populations via the GAL4-UAS system. However, transgenic expression typically labels the majority of the targeted neuronal type, often distributed across multiple brain regions. Viral delivery can provide regional specificity, but achieving precise targeting in the compact larval zebrafish brain remains technically challenging. Our method employs the transient transgene technique in zebrafish to express helper proteins. This involves temporarily introducing GAL4-UAS binary DNA plasmids into the fish’s genome through microinjection into fertilized eggs (see Figures 1A and 2A). It enables sparse expression of TVA and G in genetically defined cell types, even in a single neuron. Tracing from a single neuron can help cell-subtype-specific circuit dissection.

Furthermore, we linked the two helper protein genes with the 2A element to co-express TVA and G in the same neuron (see Figures 1A and 2A). Co-expression guarantees that the tracing is monosynaptically restricted. If G is expressed in neurons lacking TVA and a fluorescent indicator, these cells may act as input cells for the initially infected starter cells, resulting in further trans-complementation and spread of RV*d*G across the second and even more synaptic steps (Callaway and Luo, 2015). Previous studies expressed TVA and G separately, either using two transgenic lines or dual viral vectors (Dohaku et al., 2019; Satou et al., 2022), raising the risk of their expression in different neurons and potentially compromising monosynaptic specificity.

Our study demonstrated that the transient expression of helper proteins effectively facilitates subsequent viral infection and trans-complementation, offering a time-efficient and cost-effective alternative to generating stable transgenic lines. It also addresses the potential cytotoxicity effects associated with the use of helper viral vectors. With commercially available recombinant RV tools, our method can be easily implemented by zebrafish research laboratories using three commonly used techniques: plasmid construction, microinjection (with DNA constructs and recombinant RV injected 3–6 days apart), and live optical imaging.

### Key factors that influence tracing efficiency

To enhance RV*d*G tracing efficiency, we introduced the CVS-N2c rabies strain in zebrafish for the first time. It has been reported that the CVS-N2c strain exhibits significantly higher tracing efficiency in mice compared to the SAD-B19 strain, with an improvement of almost tenfold (Reardon et al., 2016). In zebrafish larvae, CVS*d*G, when trans-complemented with its native N2cG, also showed an average increase close to tenfold compared to SAD*d*G trans-complemented with its native B19G (see Figure 2D and Figure 1,2—table supplement 1, CI_neuron_). This improvement may be attributed to the reduced replication and protein expression of the CVS-N2c (Reardon et al., 2016). This could result in decreased toxicity to host cells, longer reproductive times, and ultimately higher spread efficiency. Consistently, in zebrafish larvae, we found that neurons infected with CVS-N2c displayed lower fluorescence intensity and longer survival times compared to those infected with SAD-B19. Notably, N2cG exhibited higher tracing efficiency than B19G when they were used to trans-complement the same CVS*d*G (see Figure 2D and Figure 1,2—table supplement 1). This could be attributed to the greater neurotropism of N2cG compared to B19G, or potential mismatches between the cytoplasmic domain of B19G and the CVS-N2c capsid. Further investigation into the impact of different G proteins on RV*d*G tracing efficiency in zebrafish would benefit from the construction of chimeric G proteins for comparison (Kim et al., 2016).

Besides the virus strain, temperature also plays a pivotal role. Under the optimal temperature for virus replication at 36 °C, the tracing efficiency of CVS-N2c increased more than tenfold in comparison with that under the standard rearing temperature for zebrafish at 28 °C. This effect was more pronounced for SAD-B19, showing an increase of over 80 times (see Figure 2D and Figure 1,2—table supplement 1). Importantly, no significant increase in mortality or behavioral abnormalities was observed in either non-injected or virus-injected larvae maintained at elevated temperatures. In virus-injected (CVS-N2c) larvae, survival rates decreased slightly from 100% to ∼80% at both 28 °C and 36 °C, likely due to electrode insertion-related injury rather than temperature-dependent toxicity. Virus-injected larvae could be maintained at 36 °C for over a month without adverse consequences, indicating that elevated temperature is well tolerated. This is consistent with a previous report showing no temperature-dependent effects on various behaviors in adult zebrafish between 28 °C and 36 °C (Satou et al., 2022).

The expression level of G proteins is another determinant for efficient rabies tracing (Kim et al., 2016). Building upon the codon optimization of the G protein, we further increased the expression of N2cG by inserting the Tetoff element upstream of the N2cG gene (A-N2cG) in the UAS helper plasmid (see Figure 2-figure supplement 1F, Figure 2D, and Figure 1,2—table supplement 1). This integration enables dual amplification of N2cG and TVA expression by both the GAL4-UAS and tTA-TRE binary systems. Combining the three key optimal conditions: CVS-N2c strain, 36 °C temperature, and A-N2cG, we have achieved the highest tracing efficiency in zebrafish to date, with a 20-fold increase compared to previous reports (Satou et al., 2022).

To assess the efficacy of our trans-synaptic labeling relative to endogenous connectivity, we quantified PC, GC, and IOC populations based on transgenic labeling in *Tg(2×en.cpce-E1B:tdTomato-CAAX)*, *Tg(cbln12:GAL4FF);Tg(5×UAS:EGFP)*, and *Tg(elavl3:H2B-GCaMP6s)* lines, respectively. At ∼7 dpf, we counted approximately 330 PCs, 2300 GCs, and 330 IOCs in fish brain. Considering developmental scaling and anatomical constraints (e.g., a subset of GCs project exclusively ipsilaterally), we estimate that by 10–14 dpf, a single PC receives ∼1000–2000 GC inputs. The IOC-PC connectivity is primarily one-to-one (Hamling et al., 2015; Hsieh et al., 2014). Under optimal tracing conditions, we labeled an average of ∼20 GC inputs per PC (∼1–2% capture rate) and observed IOC labeling in 7 of 248 starter PCs (∼3%). While the capture efficiency is modest, it aligns with values reported in mammalian systems (Callaway and Luo, 2015; Wall et al., 2010). Nevertheless, this indicates inherent limitations of rabies virus-based circuit tracing in zebrafish. Fundamentally addressing this limitation may require the development of viral tracing tools engineered from viruses that naturally infect fish (Lei et al., 2022).

### Monosynaptic specificity of the tracing

Employing the cerebellar circuit as a model, we examined the complete pattern of traced neurons from PCs in larval zebrafish and found that CVS*d*G was able to trace well-known presynaptic neurons of PCs, including both intra-cerebellar GCs and extra-cerebellar IOCs (see Figure 2G–I). Importantly, IOCs were the only labeled cells outside of the cerebellum, even after extended periods allowing for viral spread, demonstrating the retrograde and monosynaptic specificity of the rabies viral tracer in the zebrafish brain.

We further provided functional evidence that CVS*d*G, even at its highest efficiency with A-N2cG, spreads retrogradely between neurons through synaptic connections (see Figure 2J–M). In all experiments involving the successful activation of PCs in response to electrical stimulation of single GCs, we consistently did not observe PC activation after ablating the GCs. In addition, as shown in Figure 2L, calcium imaging at 5 Hz revealed PC responses with an average latency of 152 ± 35 ms (mean ± SEM), indicative of immediate and likely monosynaptic activation. Together with the anatomical tracing data, these findings provide strong evidence that CVS*d*G spreads retrogradely and monosynaptically from postsynaptic PCs to functionally connected presynaptic GCs.

### Low cytotoxicity that enables functional experiments in a wide time window

We found that the CVS-N2c strain did not significantly affect the neuronal health and physiology of starter cells for at least 10 days after infection (see Figure 3D–L). Meanwhile, consistent with previous reports in mice (Reardon et al., 2016), the SAD-B19 strain exhibited higher cytotoxicity than CVS-N2c, resulting in more cell death of starter cells (see Figure 3D). It indicates that CVS-N2c is more suitable for application in zebrafish. We could observe the labeling of starter cells even one month after CVS-N2c infection, and the infected zebrafish could be raised to adulthood.

A potential concern is whether viral titer or injection volume contributes to cytotoxicity. However, RV*d*G-EnvA infection generally occurs at a rate of one viral particle per cell (Zhang et al., 2024), indicating that a higher viral dose does not proportionally increase the viral load per infected cell, and is therefore unlikely to enhance tracing efficiency or aggravate cytotoxicity at the single-cell level. We used the highest viral titer that could be reliably generated in our system (see Materials and methods) and tested an increased injection volume of 20 nL (i.e., 4–10 times the standard injection volume). Injection at this volume resulted in deformation of the larval brain and was therefore not used in subsequent tracing experiments, while we observed no detectable reduction in starter cell survival within the experimental timeframe, supporting the notion that viral cytotoxicity is primarily driven by viral replication rather than by viral dose.

The enhanced G protein expression achieved through the introduction of the Tetoff element (i.e., A-N2cG) in the helper plasmid improved tracing efficiency, but also resulted in elevated levels of cytotoxicity, though lower than that induced by SAD-B19 (see Figure 3D). This indicates an association between the expression levels of G proteins and cytotoxicity (Faber et al., 2002; Ohara et al., 2013). Given the transferred RV is deficient in G, it is expected that retrogradely infected input cells will have lower cytotoxicity levels than starter cells. Therefore, input cells will survive for longer periods of time.

Based on our experience, the CVS-N2c can be microinjected into zebrafish larvae between 3 and 6 dpf, and stable and viable labeling of traced circuits can be observed within 6 to 16 dpi. Therefore, the observation time window ranges from 9 to 22 dpf, spanning the larval ages commonly used for whole-brain functional imaging studies (Cong et al., 2017; Shang et al., 2024).

### Transneuronal labeling of glial cells

Transneuronal labeling of radial glia mediated by VSV, a member of the Rhabdoviridae family like RV, has been commonly observed in larval zebrafish (Kler et al., 2021; Ma et al., 2020). In our experiments, labeled radial glia were frequently located near the somata and dendrites of starter neurons (see Figure 1D and Figure 1—figure supplement 2). In zebrafish, radial glia—also referred to as radial astroglia—are considered functional equivalents of mammalian astrocytes. These cells are known to participate in tripartite synapses and may express endogenous receptors capable of mediating RV*d*G uptake at postsynaptic sites. This anatomical proximity suggests that rabies-based trans-synaptic tracing could not only resolve neuronal connectivity but also identify glial populations that are potentially synaptically associated with and functionally coupled to defined neuronal targets. Leveraging this feature offers a promising strategy for investigating neuron-glia interactions in vivo.

Glial labeling typically emerged around 2 dpi and increased over time; however, the rate of labeling plateaued or declined during the late phase (6–10 dpi), in contrast to the continued increase in neuronal labeling (see Figure 2—figure supplement 1C and G, and Figure 2—figure supplement 2). We found that glial cells labeled at different early time points showed a convergent loss of fluorescence during the same later time window (see Figure 2—figure supplement 3). This age-dependent decline, rather than duration-dependent, is consistent with a model of abortive infection, in which the host’s innate antiviral responses suppress viral replication, preventing progeny virion production and resulting in the eventual loss of reporter expression (Tian et al., 2018). In accordance, we did not observe punctate fluorescent debris characteristic of apoptotic cell death. This suggests that the loss of fluorescence reflects functional silencing of viral gene expression rather than glial degeneration. These findings imply that abortive infection becomes more prominent with larval maturation, potentially accounting for the negligible glial labeling observed in adult zebrafish (Dohaku et al., 2019; Satou et al., 2022).

Notably, the eventual disappearance of glial fluorescence does not indicate a lack of initial viral entry. Glial infection likely competes with neuronal infection for viral particles, thereby reducing neuronal tracing efficiency. Future studies should elucidate the molecular mechanisms underlying glial susceptibility, particularly receptor-mediated viral entry at synaptic sites. Targeted suppression of viral receptors in glial cells may help minimize off-target labeling and enhance the efficiency of neuronal circuit tracing.

### Applicability and future extension of the tracing strategy

Owing to the compact small brain of larval zebrafish, the interpretability of rabies-based trans-monosynaptic tracing critically depends on the configuration of starter cells. We emphasize that this tracing strategy is readily interpretable when starter cells are sparse, limited to single neurons, or restricted to a defined cell-type population. Under these conditions, the identity and circuit order of labeled presynaptic neurons can be inferred with high confidence.

When TVA and G proteins are broadly expressed, interpretation becomes inherently ambiguous. Starter neurons comprising multiple neuronal types may be infected over overlapping time windows, preventing newly labeled cells from being unambiguously assigned to a specific type of starter neurons. Moreover, if newly labeled cells also express G proteins, secondary or multi-step propagation may occur, complicating the interpretation of circuit order. Variability in projection distance and synaptic strength may bias transsynaptic spread timing, further confounding the assignment of wiring relationships even when high-temporal-resolution imaging is available.

In zebrafish, sparse or controlled starter configurations can be achieved through genetically-defined transient transgene expression, as used in this study, as well as through single-cell electroporation or light-induced, single-cell-controlled (Tao et al., 2025) expression of helper proteins. Looking ahead, strategies that enable tighter spatiotemporal control of glycoprotein expression, such as transgenic lines with light-inducible G protein expression, will extend rabies-based tracing toward reliable analysis of multi-step or densely interconnected circuits in individual live larval zebrafish using long-term time-lapse imaging.

### Functions and expandability of the transgenic framework

In the transgenic reporter framework (i.e., *elavl3DoDioBR*) (see Figure 4A), we designed three expansion modules: 1) a promoter (*elavl3*) module that allows genetic access to the expression of reporter cassettes in specific cell types (e.g., *vglut2a* and *gad1b* gene promoters for excitatory and inhibitory targeting, respectively); 2) a Cre-dependent “Switch-On” reporter (tdTomato-CAAX) module that enables the expression of structural and functional tools in traced neurons following the spread of Cre-expressing virus, such as photoconvertible fluorescent proteins and optogenetic tools; 3) a Cre-dependent “Switch-Off” reporter (H2B-mTagBFP2) module that allows a default reporter to be silenced in infected neurons. In addition, the helper plasmid can serve as an additional expansion module that works together with the Cre-dependent transgenic reporter, allowing for full genetic visualization and dissection of neural circuits (Luo et al., 2018).

In this study, we used the *elavl3* promoter to restrict the fluorescent reporter to neurons and the “Switch-On” reporter tdTomato-CAAX to label the membrane of traced neurons. This enables clear neuron morphology labeling (see Figure 4C), facilitating the reconstruction of traced neural circuits based on single-neuron morphology. We used H2B-mTagBFP2 as the default “Switch-Off” reporter to provide a reference expression pattern for registering and integrating circuit-tracing images from different individual fish and experiments. As an application demonstration, we applied the transgenic framework to map the monosynaptic inputs to PCs from GCs in the cerebellum of larval zebrafish at ∼2.5 weeks old (see Figure 4). Interestedly, we uncovered two properties of the cerebellar circuit: 1) the ipsilateral preference of GC-to-PC connections; 2) the subtype specificity of GC-to-PC connections. Further investigation is required to elucidate the functional implications of subtype-specific GC-to-PC connections and the apparent contradiction between the bilateral projection of GC parallel fibers and the GCs’ preference for ipsilateral connections to PCs. The reference expression pattern can serve as a reference atlas to integrate other modal data from brain-wide images with information on gene expression, neurochemistry and neuronal activity (Wang et al., 2025). This will largely enhance our understanding of neural wiring patterns in relation to other neural characteristics (Leergaard and Bjaalie, 2022).

## Conclusion

Our study established an applicable and effective viral circuit tracing system for larval zebrafish. We believe that this tracing system will offer an avenue for the integrated structural and functional dissection of neural circuits underlying a wide range of brain functions in larval zebrafish.

## Materials and methods

### Animals

Adult zebrafish (*Danio rerio*) were reared under standard laboratory conditions, with temperatures set at 28 °C, and a light/dark cycle of 14 hr/10 hr. The larval zebrafish used were in *nacre* background (transparent strain with no melanophore pigmentation) and at 3–21 dpf. The sex of the larvae at this stage is not determined. All experimental procedures were approved by the Animal Care and Use Committee of the Center for Excellence in Brain Science and Intelligence Technology (CEBSIT), Chinese Academy of Sciences (NA-046-2023).

### Generation of DNA constructs and transgenic lines

The GAL4-UAS binary plasmids were applied to express fluorescent protein, TVA protein, and viral G in specific neuronal types. To randomly target neurons, we used *elavl3:GAL4-VP16* (Du et al., 2018) or *vglut2a:GAL4FF* (Du et al., 2025). To target Purkinje cells (PCs), we generated a construct to drive the *GAL4FF* under the control of *cpce* (*ca8* promoter derived PC-specific enhancer element) fused to *E1B* (Namikawa et al., 2019). The *cpce* sequence was PCR-amplified from the zebrafish genome DNA, and the *cpce-E1B:GAL4FF* was flanked by a minimal *Tol2* transposase recognition sites (Kawakami, 2005). We generated seven helper plasmids (abbreviated names are in parentheses): *14×UAS-E1B:EGFP-P2A-TVA* (*UGT*), *14×UAS-E1B:EGFP-P2A-TVA-T2A-B19G* (*UGTB*), *14×UAS-E1B/5×UAS-hsp:EGFP-P2A-B19G-T2A-TVA* (*UGBT*, the *14×* and *5×UGBT* were used with *elavl3:GAL4-VP16* and *cpce-E1B:GAL4FF*, respectively), *5×UAS-hsp:sfGFP-P2A-N2cG-P2A-TVA* (*UGNT*), *5×UAS-hsp:sfGFP-P2A-Tetoff-N2cG-P2A-TVA* (*UGNT-A*), and *5×UAS-hsp:GCaMP6s-P2A-Tetoff-N2cG-P2A-TVA* (*UG6sNT-A*). The *sfGFP*, *N2cG*, *B19G*, *TVA*, and *Tetoff* were codon optimized for zebrafish and *de novo* synthesized; the *GCaMP6s* was PCR-amplified from *elavl3:GCaMP6s* plasmid (gift from M. B. Ahrens, Janelia Research Campus, USA). All these elements were sequentially assembled with 2A sequences and the linearized *miniTol2-UAS* backbone to generate polycistronic vectors through In-Fusion cloning. Sparse expression of helper proteins was achieved by the microinjection of a mixture (1 nl) of *elavl3* or *cpce*-driven GAL4 plasmid, UAS-driven helper plasmid, and *Tol2* transposase mRNA into one-cell stage zebrafish embryos at a concentration of 30 (10+10+10) ng/μl using an air-puffed pressure injector (MPPI-3, ASI).

To generate *Tg(2×en.cpce-E1B:tdTomato-CAAX)*, we created a construct to drive *tdTomato* with a 3’ membrane tag encoding the *CAAX* box of human Harvey Ras (*CTGAACCCTCCTGATGAGAGTGGCCCCGGCTGCATGAGCTGCAAGTGTGTGCTCTCC*) under the control of two tandem *cpce* fused to E1B. To generate *Tg(cbln12:GAL4FF)* for labeling granule cells (GCs) in the cerebellum, we amplified a ∼2-kbp regulatory element from zebrafish genome DNA by PCR for the *cbln12* (*cerebellin12*) gene and subcloned it into the linearized *miniTol2-GAL4FF* backbone. To generate *elavl3DoDioBR*, we first synthesized a zebrafish codon-optimized *mTagBFP2 de novo* and then constructed an intermediate plasmid containing *Tetoff-DO_DIO-Hsa.H2B-mTagBFP2_tdTomato-CAAX* using In-Fusion cloning. Subsequently, we subcloned this target fragment into the linearized mini*Tol2-elavl3* backbone to generate the final vector. Transgenic zebrafish were prepared using the *Tol2* transposase-based approach. Purified target plasmid DNA was microinjected with the *Tol2* transposase mRNA into one-cell stage zebrafish embryos at a concentration of 60 (30+30) ng/μl. Injected embryos were raised to sexual maturity to identify founder fish. The *Tg(cbln12:GAL4FF)* was crossed with *Tg(5×UAS:EGFP)* to visualize GCs.

### Production of EnvA-pseudotyped rabies virus

The EnvA-pseudotyped rabies viruses (RV), including SAD*d*G-mCherry[EnvA], CVS*d*G-tdTomato[EnvA], and CVS*d*G-mCherry-2A-Cre[EnvA], were rescued and prepared following the previously reported method (Lin et al., 2022; Osakada and Callaway, 2013). Rabies vectors were prepared in phosphate-buffered saline (PBS) and stored at –80 °C in aliquots for experimental use. The injected titers were 2 × 10^8^ infectious units per ml (IU/ml) for SAD*d*G-mCherry[EnvA] and CVS*d*G-tdTomato[EnvA], and 3 × 10^8^ IU/ml for CVS*d*G-mCherry-2A-Cre[EnvA]. We found that mCherry fluorescence after CVS*d*G-Cre[EnvA] infection in vivo was much weaker than that of transgenetically expressed tdTomato-CAAX, and thus did not affect the observation and reconstruction of neuronal morphology.

### Virus application

At 3–6 dpf, larvae expressing helper proteins in specific neurons were anesthetized using 0.03% tricaine methanesulfonate (MS-222) and mounted laterally in 1.0% low-melting agarose (Sigma) on a glass-bottom dish filled with the extracellular solution consisted of (in mM): 134 NaCl, 2.9 KCl, 2.1 CaCl_2_•2H_2_O, 1.2 MgCl_2_•6H_2_O, 10 HEPES, and 10 glucose (pH = 7.8, osmolality = 290LmOsm). Virus delivery was performed under a standard high-magnification upright (135×, Nikon SMZ18) or inverted fluorescence microscope (200×, Olympus IX51), enabling clear visualization of target neurons labeled with sfGFP or GCaMP6s. Approximately 2–5 nl of virus suspension was injected adjacent to the cells of interest in one hemisphere using a glass micropipette with a ∼10 µm tip diameter. The micropipette was held by a microinjector (Nanoliter2010, WPI) and positioned using a motorized manipulator (MC1000e, Siskiyou) to ensure accurate placement. Injection into the brain tissue but not ventricle was necessary to allow the virus solution to disperse locally and reach surrounding target neurons for efficient infection. After injection, the fish were allowed to rest for one minute before withdrawing the pipette. Then, they were placed in standard tanks in an incubator set at temperatures of 28 °C or 36 °C.

### Confocal and two-photon imaging

Larval fish were mounted dorsal-side up in 1.5% low-melting agarose on a custom-made chamber. Structural live imaging of virus-injected fish and transgenic lines was conducted using an upright confocal laser scanning microscope (FV3000, Olympus) equipped with a 20× water-immersion objective (N.A., 0.9). Image acquisition was performed at a lateral resolution of 0.5 × 0.5 μm and a *z*-step size of 1 μm. For virus-injected fish, the imaging dimension was based on the specific region of viral infection and spread. For transgenic lines, the entire brain was imaged by multiple volumes, which were then stitched together using a custom ImageJ (http://fiji.sc) macro based on the Grid/Collection stitching plugin (Preibisch et al., 2009). Functional Ca^2+^ imaging of PCs expressing GCaMP6s was performed using an upright two-photon microscope (FVMPE-RS, Olympus) equipped with a 25× water-immersion objective (N.A., 1.05) at a wavelength of 920 nm. The calcium activity was recorded at ∼5 Hz on single optical plane.

### Electrical stimulation of single granule cell

Within 6 to 10 days after virus injection, larvae were paralyzed with pancuronium dibromide (PCD, 1mM, Tocris 0693) for 1 min and then mounted in 1.5% low-melting agarose on a chamber filled with the extracellular solution. The agarose and skin over the caudal side of the cerebellum were removed for the access of stimulation electrodes. Stimulation microelectrodes (1–2 µm tip opening) were created using borosilicate theta glass capillaries (BT-150-10, Sutter Instrument) through a micropipette puller (P-97, Sutter Instrument). Then a silver wire was inserted into the electrode filled with the extracellular solution. The electrode was positioned at and attached to the cell body of the target GC, which was identified by the fluorescence and morphology using confocal imaging. The stimulation was delivered by a flexible stimulus isolator (ISO-Flex, AMPI), consisting of 5 pulse trains (6–8 V, 10 ms duration for one pulse, 10 pulses, 50 Hz) with inter-trial intervals of 40 s. The electrical stimulation of single GC was synchronized with the Ca^2+^ imaging of PCs by a stimulator (Master-8, AMPI).

### Two-photon laser-based ablation of granule cells

A two-photon laser (795 nm) was used to ablate the GC after electrical stimulation. Confirmation of a successful ablation involved the identification of a bulb-like structure in the bright-field image and the absence of the tdTomato fluorescence signal from the targeted GC body. The efficiency of the ablation was further validated one hour later, before performing post hoc electrical stimulation at the same position using the bulb-like structure and adjacent fluorescent cells as references.

### In vivo whole-cell recording of Purkinje cells

Within 6 to 15 days after virus injection, larvae were paralyzed with α-bungarotoxin (1mM) for 1 min, then mounted in 1.5% low-melting agarose on a chamber filled with the extracellular solution. The agarose and skin over the caudal side of the cerebellum were removed for the access of recording pipettes. A recording micropipette (15–20 MΩ, 1–1.5 µm tip opening) filled with internal solution was approached to the target PC by a motorized micromanipulator (MP-225, Sutter Instruments). The internal solution consisted of (in mM): 100 K-gluconate, 10 KCl, 2 CaCl_2_•2H_2_O, 2 Mg_2_•ATP, 0.3 GTP•Na_4_, 2 phosphocreatine, 10 EGTA, and 10 HEPES (pH = 7.4, osmolality = 270LmOsm). The target PC was identified by double fluorescence (sfGFP^+^ and tdTomato^+^) using confocal imaging and was recorded at whole-cell mode with similar procedures as our previous work (Mu et al., 2012). The EPSP recording and current-step delivery were conducted in current-clamp mode, and the EPSC was assessed in voltage-clamp mode. Data were acquired using a patch-clamp amplifier (MultiClamp 700B, Axon Instruments) and a digitizer (Digidata 1440A, Axon Instruments), and signals were sampled at 10 kHz with Clampex 10.6 software (Molecular Devices).

### Visual stimulation

Larvae examined were not anesthetized and were fully embedded in low-melting agarose. Visual stimuli were generated by a custom program written in Python and presented using an LCD screen covered with red filter paper to avoid spectrum interference. The visual stimuli basically followed a prior study (Knogler et al., 2019). Briefly, three trials were presented and each trial consisted of three types of stimuli in the following order: (1) four-directional whole-field gratings in black and white (including three forward, one backward, one leftward, and one rightward directions); (2) windmill patterns in black and white rotating at 0.2 Hz and varying in velocity following a sine function (including six whole, two right-half, and two left-half filed stimuli presented in alternating clockwise and counterclockwise directions); (3) whole-field flashes of 1-second duration (alternating between white and black, repeated 4 times). Except for the flash, each stimulus lasted 5 seconds, with a 5-second inter-stimulus interval.

### Quantification of RV*d*G[EnvA]-based circuit tracing

Cells were counted by manually annotating the center of their cell bodies in image stacks using the Cell Counter plugin in ImageJ. This was done for cells showing green (sfGFP/GCaMP6s; i.e., infected and uninfected TVA-expressing cells), red (tdTomato/mCherry; i.e., traced input cells and starter cells), or yellow (both green and red; i.e., initially infected starter cells) fluorescence separately. To count the traced neurons and glia separately, we also manually identified and marked the neurons based on their significant morphological distinctions with glia in larval zebrafish. The number of traced glia was then calculated by subtracting the number of starter cells and traced neurons from the total number of cells with red fluorescence. To determine virus tracing efficiency, a convergence index (CI) was calculated as the number of traced inputs (neurons, glia, or all) divided by the number of starter cells.

### Analysis of calcium imaging data

The acquired time-lapse images were aligned first, and individual regions of interest (ROIs) were manually circled on the averaged image. Each ROI corresponds to either a starter cell or a non-starter cell, and the mean grayscale value within each ROI over time was represented as F. In the electrical stimulation experiment, the baseline value F_0_ was calculated as the average value of F from a 10-second interval before each stimulus. In the visual stimulation experiment, F_0_ was calculated as the average value of F over a 30-second period preceding (F − F_0_)/F_0_ the stimuli. The relative change in the intensity of each ROI was calculated using the formula Response latency was defined as the time interval between the onset of electrical stimulation and the first post-stimulus imaging frame in which ΔF/F_0_ exceeded the pre-stimulus baseline by more than 3× standard deviations (SD). The area under the curve (AUC) and the number of Ca^2+^ activity events were determined by applying a threshold of 2× SD of a 60-second baseline activity. All analyses were performed with custom Python scripts.

### Template generation and registration

We first generated two two-channel templates (17 dpf), *Tg(2×en.cpce:tdTomato-CAAX);elavl3DoDioBR* and *Tg(cbln12:GAL4FF);Tg(5×UAS:EGFP);elavl3DoDioBR*, by running a shell script antsMultivariateTemplateConstruction2.sh (-d 3 -b 1 -g 0.1 -i 10 -c 2 -j 6 -k 2 -w 1x1 -f 12x8x4x2 -s 4x3x2x1vox -q 400x200x200x20 -r 1 -n 0 -m CC -t SyN ${image-inputs}) on the CEBSIT’s Linux computing cluster using the open-source Advanced Normalization Tools (ANTs) software (Avants et al., 2010). Confocal images were acquired at 1 × 1 × 1 μm resolution. Both templates were shape-based average representations from 6 transgenic larvae after 7 iterations. Using normalized cross correlation (NCC) as a metric to evaluate the registration accuracy between *elavl3DoDioBR* templates and individual *elavl3DoDioBR* scans of larvae used for circuit tracing, we determined the *elavl3DoDioBR* in *Tg(2×en.cpce:tdTomato-CAAX);elavl3DoDioBR* template space as the common coordinate reference for registration. The expression pattern of *Tg(cbln12:GAL4FF);Tg(5×UAS:EGFP)* and circuit tracing results were transformed into the common coordinate space by using *elavl3DoDioBR* as a bridge. For image registration, we used antsRegistration and antsApplyTransforms functions in ANTs, following the registration parameters for live fish scans reported in previous work (Marquart et al., 2017). Neuron reconstructions (SWC files (Cannon et al., 1998)) were aligned using antsApplyTransformsToPoints function in ANTs.

### Cerebellum delineation and neuron reconstruction

The 3D cerebellum region was delineated manually using 3D Slicer software (https://www.slicer.org/) based on the expression patterns of *elavl3DoDioBR*, *Tg(2×en.cpce:tdTomato-CAAX)*, and *Tg(cbln12:GAL4FF);Tg(5×UAS:EGFP)* in the common coordinate space. The boundary delineation was performed on *elavl3DoDioBR*, assisted by overlaying the other two patterns.

To reconstruct the morphology of neurons in virus-traced circuits, neurons expressing membrane-bound tdTomato were semi-automatically traced using the simple neurite tracer (SNT) (Arshadi et al., 2021) plugin in ImageJ. The classification of dendritic and axonal compartments of neurites was based on structural knowledge. Neurons with low fluorescence intensity and ambiguous morphology were excluded during reconstruction. The resulting individual neuron skeletons were saved in SWC format. Tracings containing undistinguished neuron skeletons were also saved as individual SWC files. Connectivity between individual starter PCs and traced GCs was determined by examining original confocal image stacks for clear appositions between GC axons and PC dendrites or somata.

The co-volume rendering of template cerebellum and cerebellum region (see Figure 4E) was performed using ParaView (Ahrens et al., 2005) (https://www.paraview.org). The co-visualization of the cerebellar region and registered neuron reconstructions (see Figure 4F,I,J) was performed using custom python scripts based on NAVis package (Schlegel et al., 2023) (https://github.com/navis-org/navis).

### Statistics

All statistical tests and graphs were performed with GraphPad Prism or Python software. Data normality was first examined with Shapiro-Wilk test. Comparisons between two groups with normal and non-normal distributions were made using two-tailed unpaired Student’s *t* test and nonparametric two-tailed Mann-Whitney test, respectively. Differences between groups involving multiple factors were analyzed using two-way ANOVA. *P* value <0.05 achieved statistical significance. Data are represented as mean ± SEM. The sample size and/or the number of replicates for each experiment are reported in the main text, figures, figure legends and/or supplementary tables.

## Acknowledgments

We are grateful to Drs. M. B. Ahrens for sharing DNA plasmids, F. Peri and K. Kawakami for sharing zebrafish lines. We thank X.L. Shen, L. Sun, S. Li, and K. Wang for their kind help on virus injection, calcium imaging, visual stimulation, and plasmid construction experiments, respectively. This work was supported by Brain Science and Brain-like Intelligence Technology - National Science and Technology Major Project (2021ZD0204500 and 2021ZD0204502), the National Natural Science Foundation of China (32321003 and 62320106010), and Shanghai Municipal Science and Technology Major Project (18JC1410100 and 2018SHZDZX05). X.F.D. is also supported by the Youth Innovation Promotion Association of CAS.

## Author contributions

X.F.D. and J.L.D. conceptualized and supervised the project. T.L.C. and Q.S.D. developed the methodology and designed the experiments with input from X.F.D. and J.L.D.. T.L.C. and Q.S.D. performed the experiments with input from X.W. and X.D.Z.. X.D.Z. created the cerebellum-related transgenic fish. K.Z.L. produced the rabies viruses under the supervision of F.Q.X.. T.L.C. analyzed data and made the illustrations with input from Q.S.D., Y.W.Z., and X.Y.N.. Y.L. helped to establish the virus injection in zebrafish. T.L.C., Q.S.D., X.F.D., and J.L.D. wrote the manuscript.

## Data availability

We have deposited the related zebrafish lines at CZRC (China Zebrafish Resource Center) and uploaded plasmid maps and sequences to Addgene. The viral vectors are available through BrainCase (Shenzhen, China). The neuron reconstructions can be downloaded at https://doi.org/10.12412/BSDC.1748509594.10001. All original code has been deposited at https://github.com/soaringdu/Proj-RVCT. Any additional information required to reanalyze the data reported in this paper is available upon request.

---

**Figure 1—figure supplement 1.**
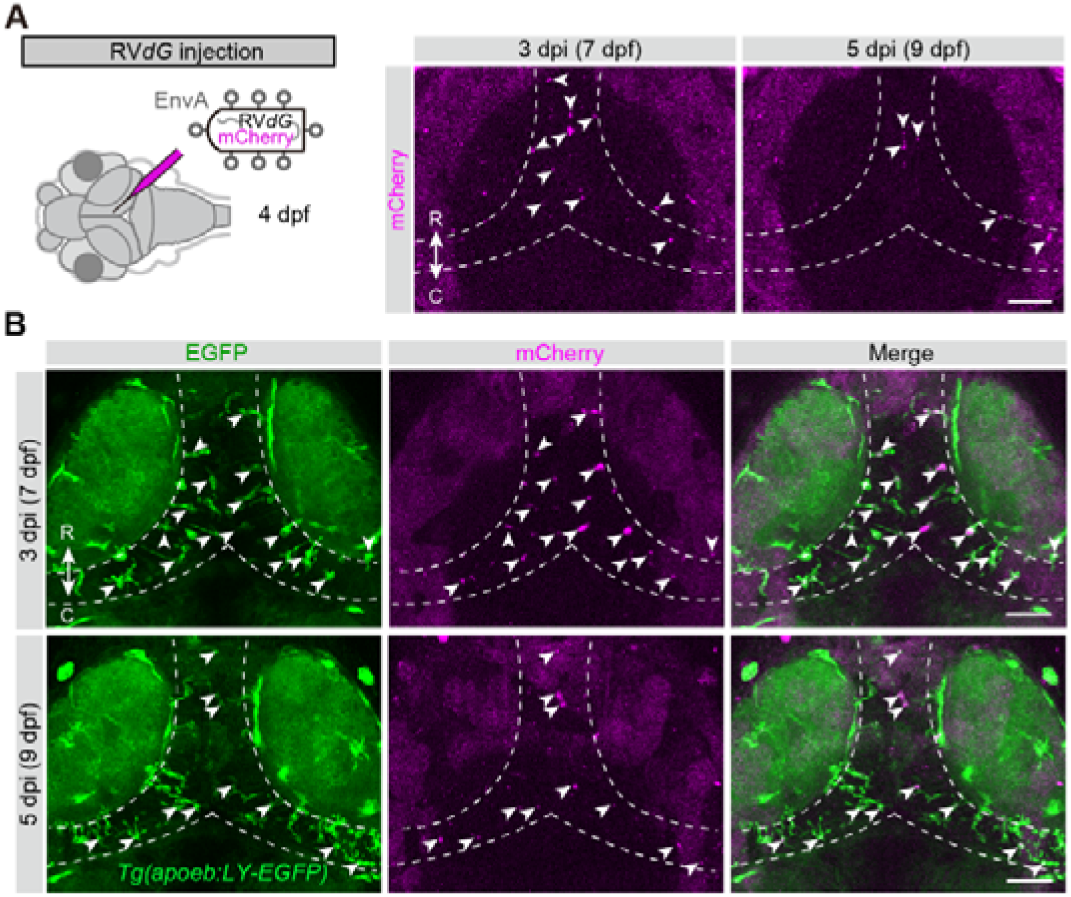
No infectiousness of RV*d*G[EnvA] in the absence of TVA. (A) Injection of SAD*d*G-mCherry[EnvA] into the brain of *nacre* larvae. Left, schematic; Right, time-lapse confocal images of the optic tectum of a WT larva injected with the virus. (B) Time-lapse confocal images of the optic tectum of a *Tg(apoeb:LY-EGFP)* larva injected with SAD*d*G-mCherry[EnvA], showing the location of mCherry-positive puncta in EGFP-positive microglia. Scale bars, 50 μm. Arrowheads, fluorescent-positive punctate structure with no discernible cellular morphology under high laser excitation; dashed lines, boundaries of the cell body layer of the optic tectum. C, caudal; R, rostral.

**Figure 1—figure supplement 2.**
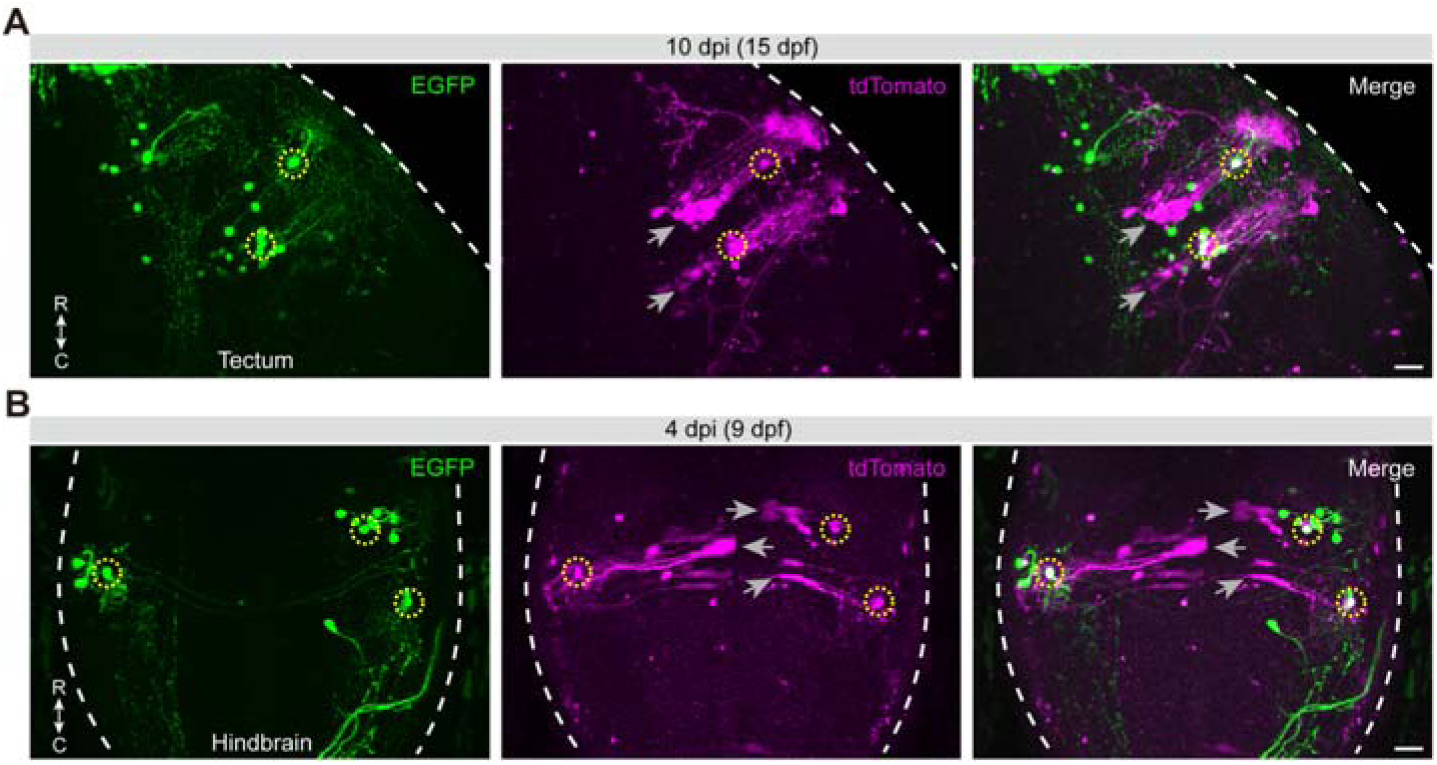
Location of transneuronally labeled glial cells. (A and B) Confocal images of the right tectum (A) and posterior hindbrain (B) from different WT larvae, showing randomly sparse *vglut2a*^+^ neurons expressing EGFP and helper proteins via *UGNT*, following infection with CVS*d*G-tdTomato[EnvA] (magenta) injected into the anterior hindbrain (see Figure 2 for the helper plasmid and virus vector). Scale bars, 20 μm. Dashed yellow circles, starter neurons (EGFP^+^/tdTomato^+^); gray arrows, transneuronally labeled radial glia (tdTomato^+^/EGFP^−^); dashed white lines, tectal or hindbrain boundaries. C, caudal; R, rostral.

**Figure 1,2—figure supplement 1.**
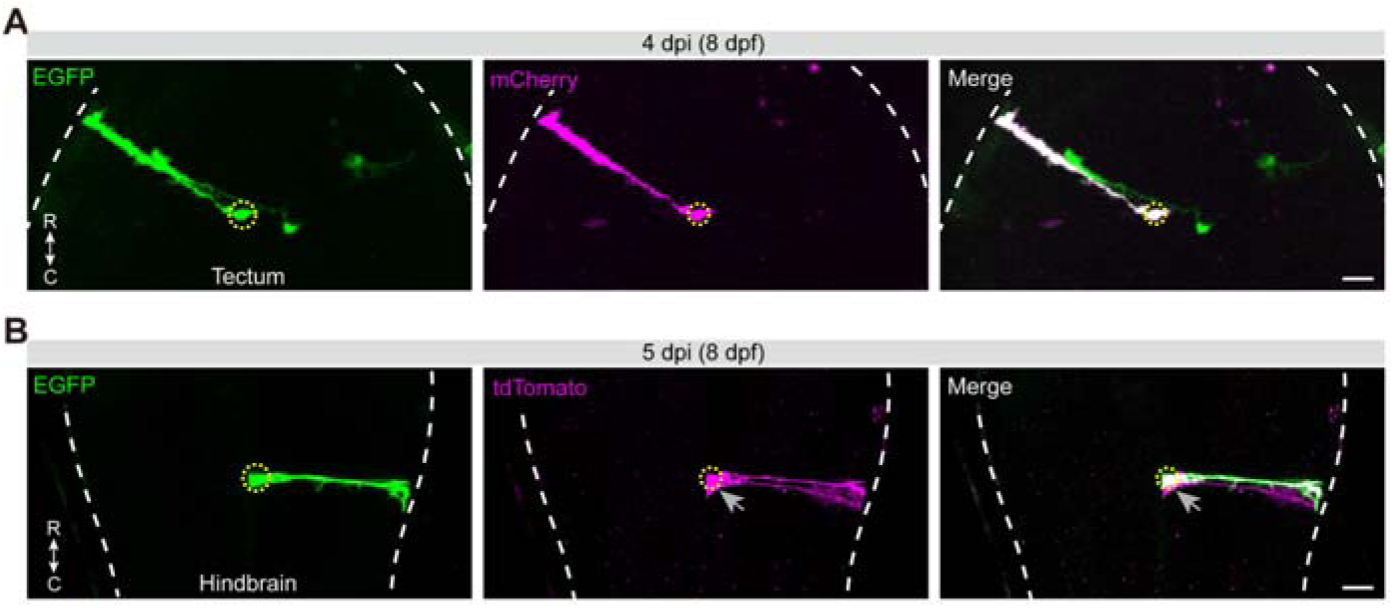
Viral tracing initiated from glia. (A) Confocal images of the tectum of a larva showing randomly sparse *gfap*^+^ glial cells expressing EGFP and helper proteins via *UGBT*, following infection with SAD*d*G-mCherry[EnvA] (magenta) injected into the anterior hindbrain at 28 °C. (B) Confocal images of the posterior hindbrain of a larva showing randomly sparse *gfap*^+^ glial cells expressing EGFP and helper proteins via *UGNT*, following infection with CVS*d*G-tdTomato[EnvA] (magenta) injected into the anterior hindbrain at 28 °C. Scale bars, 20 μm. Dashed yellow circles, starter glial cells (EGFP^+^/mCherry^+^ or EGFP^+^/tdTomato^+^); gray arrows, secondarily labeled glial cells (tdTomato^+^/EGFP^−^); dashed white lines, tectal or hindbrain boundaries. C, caudal; R, rostral.

**Figure 2—figure supplement 1.**
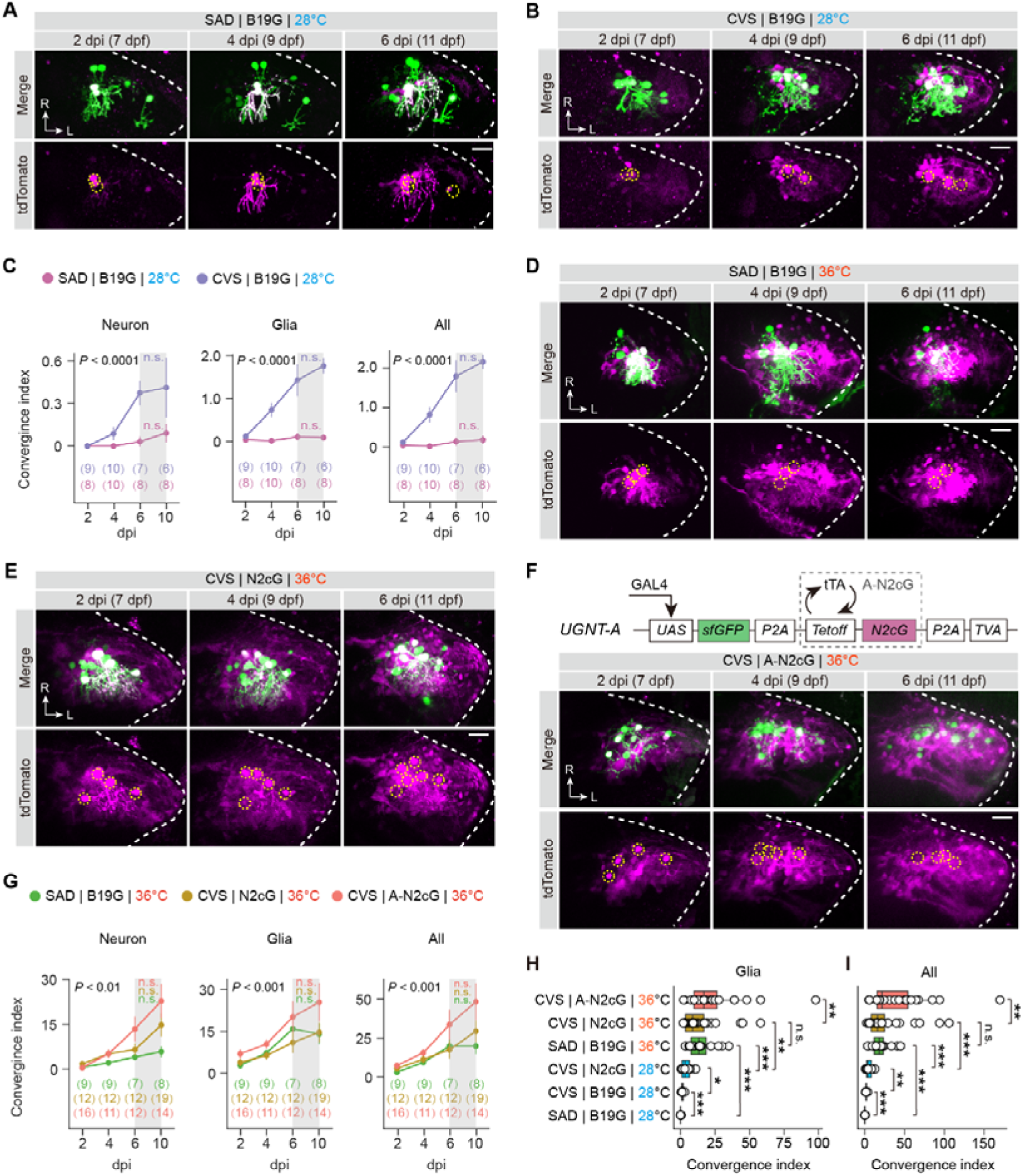
Time-lapse images of trans-complemented tracing from Purkinje cells in the cerebellum under different tracing conditions. (A, B, and D–F) Time-lapse (2, 4, and 6 dpi) confocal images of in situ complementation in Purkinje cells (PCs) under tracing conditions of SAD|B19G|28 °C (A), CVS|B19G|28 °C (B), SAD|B19G|36 °C (D), CVS|N2cG|36 °C (E), and CVS|A-N2cG|36 °C (F). The schematic drawing of the advanced helper plasmid (*UGNT-A*) is shown in (F). Dashed yellow circles, starter PCs; dashed white lines, the cerebellar boundaries. L, lateral; R, rostral. Scale bars, 20 μm. (C and G) Summary of CI for the connection of traced neurons, glia, or all cells to starter PCs at 2, 4, 6, and10 dpi under two tracing conditions at 28 °C (C) and three tracing conditions at 36 °C (G). Significant intergroup differences exist in CI for traced neurons, glia, or all cells (see the *P* values). There are no statistical differences (n.s.) between 6 and 10 dpi in any of the groups. Two-way ANOVA and nonparametric two-tailed Mann-Whitney test were used for intergroup and intragroup statistics, respectively. Error bars denote SEM. Numbers in brackets indicate the number of larvae examined. (H and I) Boxplots of CI for the connection of traced glia (H) or all cells (I) to starter PCs under six tracing conditions at 6–10 dpi. See also Figures 2C–E. Center, median; bounds of box, first and third quartiles; whiskers, minimum and maximum values. n.s., not-significant; \**P* < 0.05; \*\**P* < 0.01; \*\*\**P* < 0.001 (nonparametric two-tailed Mann-Whitney test).

**Figure 2—figure supplement 2.**
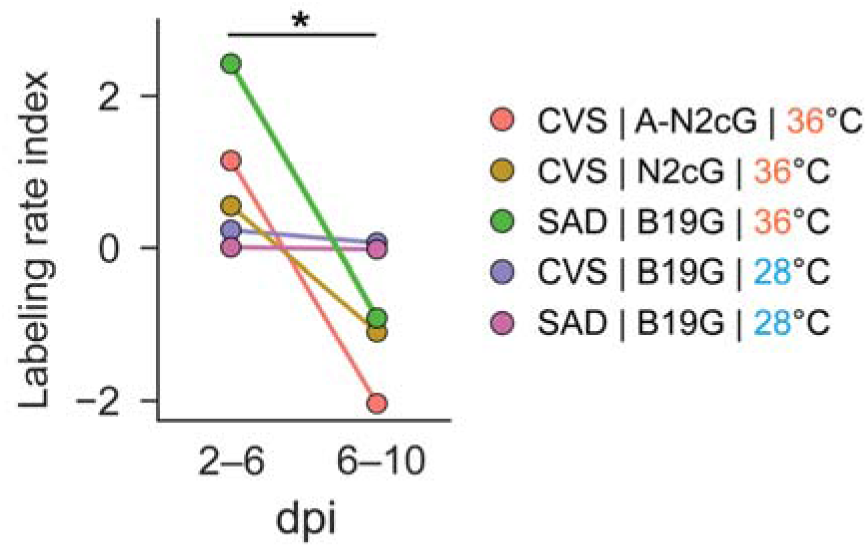
Calculation of the labeling rate index across two time intervals. The labeling rate for neurons and glial cells under each experimental condition was calculated as the daily increase in the mean CI across defined time intervals, based on data from Figure 2—figure supplement 1C and G. The mean CI was quantified separately for neurons and glia at each time point (dpi) by averaging CI values from individual fish within each group. Changes in mean CI across two consecutive time intervals (2–6 and 6–10 dpi) were normalized by the interval duration (4 days) to obtain daily labeling rates for each cell type. The labeling rate index (R_glia_ − R_neuron_) was defined as the difference between glial (R_glia_) and neuronal (R_neuron_) labeling rates. \**P* < 0.05 (nonparametric two-tailed Mann-Whitney test).

**Figure 2—figure supplement 3.**
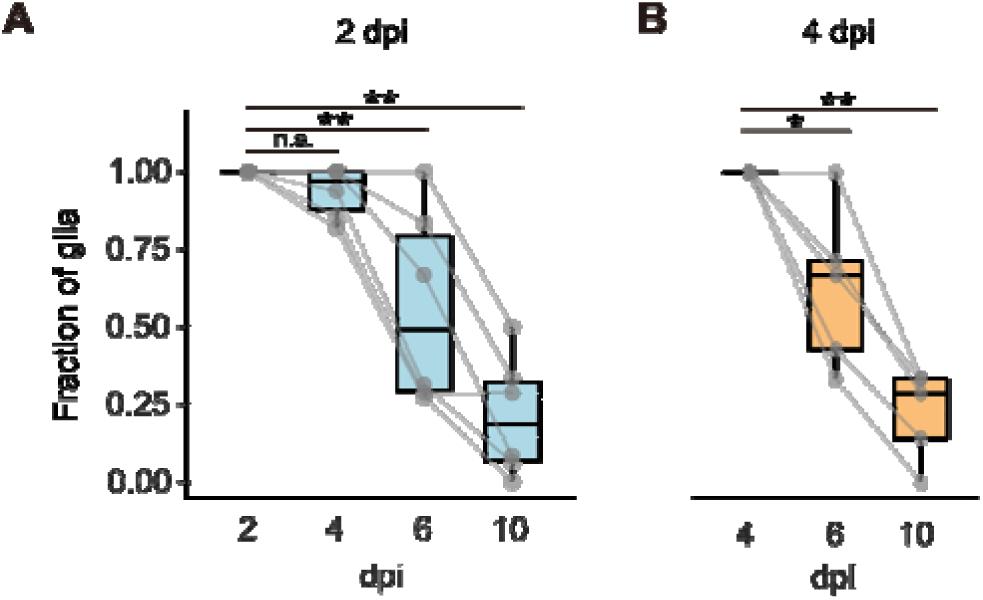
Fraction of glial cells retaining fluorescence. (A and B) Boxplots of the proportion of glial cells labeled at 2 dpi (A) or 4 dpi (B) that retained detectable fluorescence at each subsequent time point. Data are from the CVS|N2cG|36 °C group shown in Figure 2—figure supplement 1G. Box center, median; bounds of box, first and third quartiles; whiskers, minimum and maximum values. n.s., not-significant; \**P* < 0.05, \*\**P* < 0.01 (nonparametric two-tailed Mann-Whitney test).

**Figure 1,2—table supplement 1.**
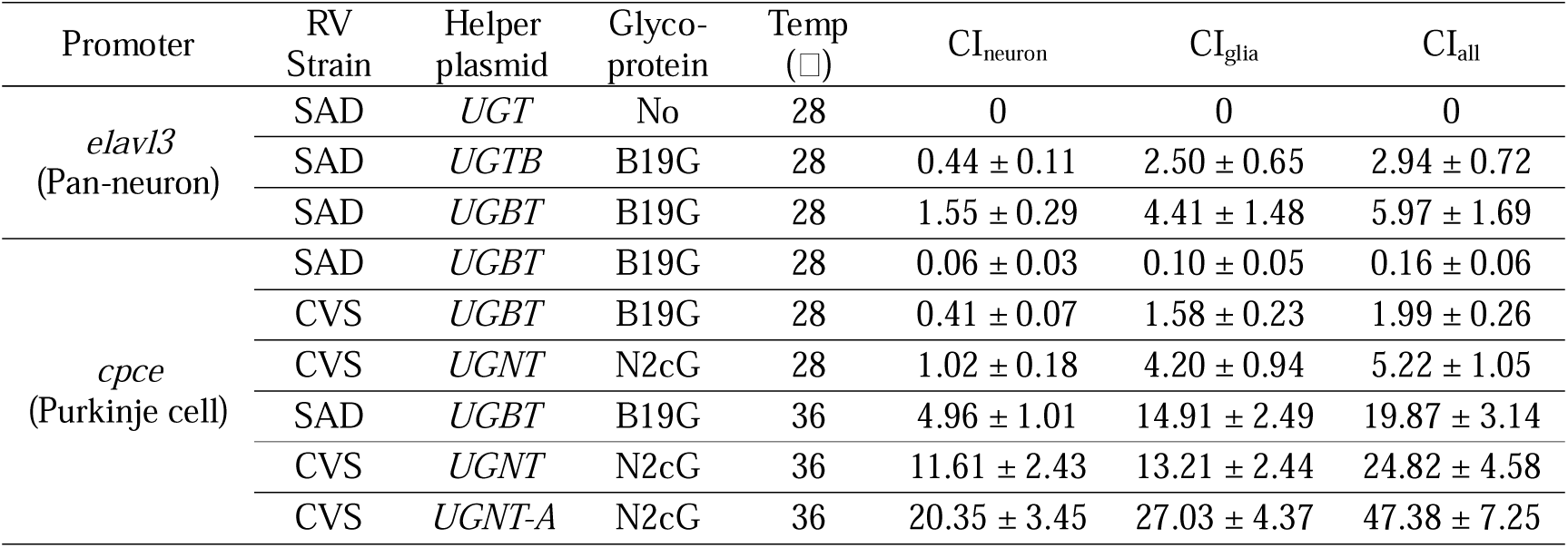
The efficiency of viral transfer under different tracing conditions in larval zebrafish. The convergence index (CI) was calculated as the number of traced cells divided by the number of starter cells.

**Figure 1,2—table supplement 2.**
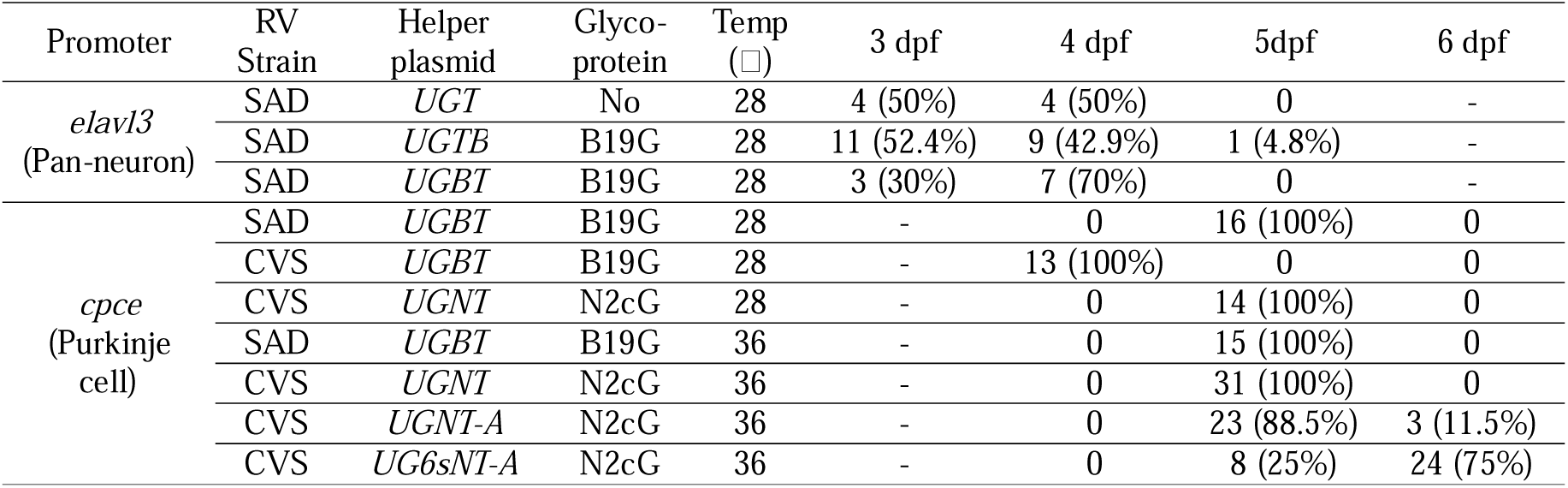
Injection age and corresponding fish numbers for each tracing condition. The proportion of fish at each injection age is shown in parentheses.

**Figure 1–4—table supplement 1.**
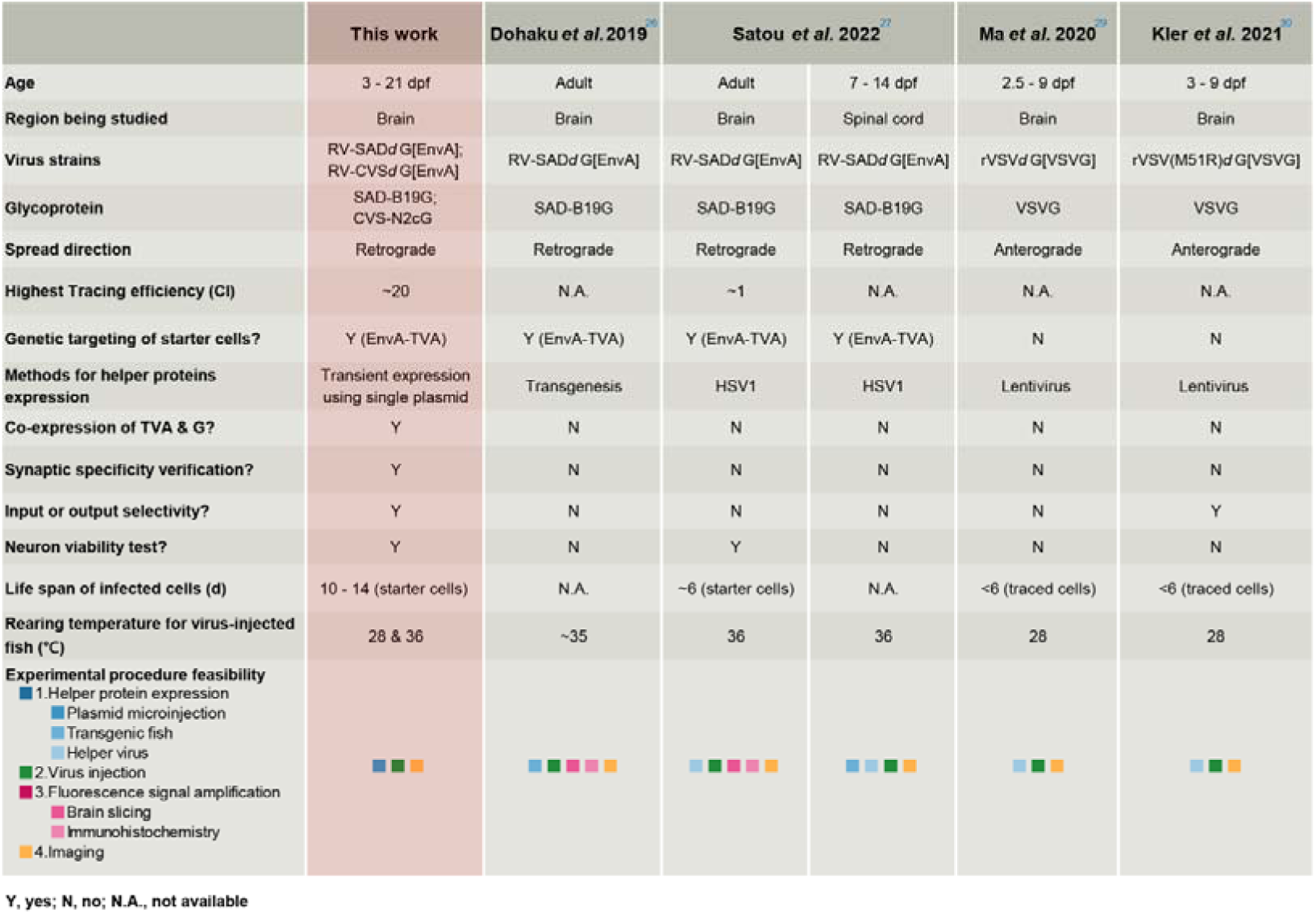
Properties of currently available viral circuit tracing systems for zebrafish.

